# Perceptual stability reflected in neuronal pattern similarities in human visual cortex

**DOI:** 10.1101/2022.08.02.502429

**Authors:** Rotem Broday-Dvir, Yitzhak Norman, Michal Harel, Ashesh D. Mehta, Rafael Malach

## Abstract

The magnitude of neuronal activation is commonly considered a critical factor for conscious perception of visual content. However, this dogma contrasts with the phenomenon of rapid adaptation, in which the magnitude of neuronal activation drops dramatically in a rapid manner, while the visual stimulus and the conscious experience it elicits remain stable. Here we report that the profiles of multi-site activation patterns and their relational geometry –i.e. the similarity distances between activation patterns, as revealed using iEEG recordings, are sustained during extended stimulation despite the major magnitude decrease. These results are compatible with the hypothesis that conscious perceptual content is associated with the neuronal pattern profiles and their similarity distances, rather than by the overall activation magnitude, in human visual cortex.

## Introduction

What is the neuronal code that underlies perceptual experiences in the human brain? Contrasting changes in conscious perceptual content while maintaining the physical stimulus conditions constant has played a central role in this quest. This approach has been employed in different tasks and conditions such as bi-stable illusions, binocular rivalry, backward masking or attentional shifts (Sheinberg and Logothetis, 1997; Tong et al., 1998; Hasson et al., 2001; Quiroga et al., 2005; Fisch et al., 2009; Davidesco et al., 2013; Gelbard-Sagiv et al., 2018). The findings from this large body of work consistently reveal that conscious perception is associated with a rapidly increased magnitude of neuronal activity, specifically in content-selective sites associated with the particular conscious experience. Similarly, comparing invisible spontaneous blinks with salient blank-screen interruptions reveals significant activation magnitude modulations associated with the conscious awareness to the interruptions in high (but not early) visual areas (Golan et al., 2016). Thus, a number of findings appear to converge in pointing to activation level as a common signature of perceptual awareness, directly linking conscious perception of sensory content with increased magnitude of neuronal activation.

However, a puzzling contradiction to these converging lines of evidence is presented by the robust and ubiquitous phenomenon of rapid visual adaptation. Rapid adaptation refers to a striking reduction in neuronal activity magnitude in the visual cortex, while the visual input is maintained constant for a prolonged duration, along with a stable internal perceptual experience of the visual content. These rapid adaptation effects appear few hundreds of milliseconds after stimulus onset (Muller et al., 1999; Kohn, 2007), and have been attributed to single-neuron response fatigue, as well as a normalization mechanism (Clifford et al., 2007; Kohn, 2007; Solomon and Kohn, 2014). These effects have been documented across the visual cortex, and were demonstrated using different measuring methods, such as fMRI, electrocorticography (ECoG), and single-cell recordings (Bandettini et al., 1997; Muller et al., 1999; Golan et al., 2016; Gerber et al., 2017; Norman et al., 2019). The neuronal response decays in high order visual areas are especially prominent, and show consistent temporal dynamics that are unaffected by the stimulus presentation duration, both in controlled laboratory picture viewing experiments (Gerber et al., 2017), as well as in more ecological, free viewing conditions (Podvalny et al., 2017). This rapid adaptation phenomenon likely shares common properties and mechanisms with the well-studied phenomena of visual adaptation and repetition suppression (Grill-Spector et al., 1999; Grill-Spector et al., 2006; Malach, 2012).

The adaptation phenomenon poses an obvious conundrum to activation magnitude-based theories of perception. Simply put, if the magnitude of neuronal activity determines perceptual awareness, how does the perception remain stable despite a massive reduction in the activation magnitude? Complementary, one may view the perceptual stability conundrum as a unique opportunity to advance our understanding of the neuronal correlates of perceptual awareness, through searching for those aspects of neuronal activity that remain stable in tandem with the stable nature of perception.

Interestingly, recently, the question of the neuronal correlates of perceptual stability has been identified as a key experimental test in a high impact adversarial collaboration. Specifically, predictions of a sustained-stimulation experiment were deemed to be diagnostic in supporting one of the two central theories of conscious experience-the Global Workspace Theory (GWT) and the Integrated Information Theory (IIT) (Melloni et al., 2021).

A third motivation to explore the adaptation phenomena relates to a recent hypothesis, inspired by classical structuralist perspectives, proposing that perceptual content is not solely dependent on the magnitude of neuronal activations, but rather on the relative similarity of its activation pattern to all other representational patterns within a specific cortical region (Malach, 2021). Interestingly, this notion has recently been extended to more global notions of cortical representations (Lau et al., 2022). In the rest of the text, we will term this pattern of similarity distances the *relational code*. Figure 1 depicts, in a schematic form, the different predictions associated with magnitude coding vs. relational coding. Thus, according to magnitude based coding, a displayed image will be perceived depending on the relevant neurons’ activity levels. In the proposed relational coding model, on the other hand, the overall magnitude of activation is not relevant, as the unique visual percept is defined by the similarity distances of its activation pattern to all other representational patterns, or, using vector space terminology-its relative location within the relevant relational geometry space. This relational code is operationally defined by calculating the correlation-based similarity distances between neuronal population activity patterns elicited by different visual stimuli within the represented category.

**Figure 1.**
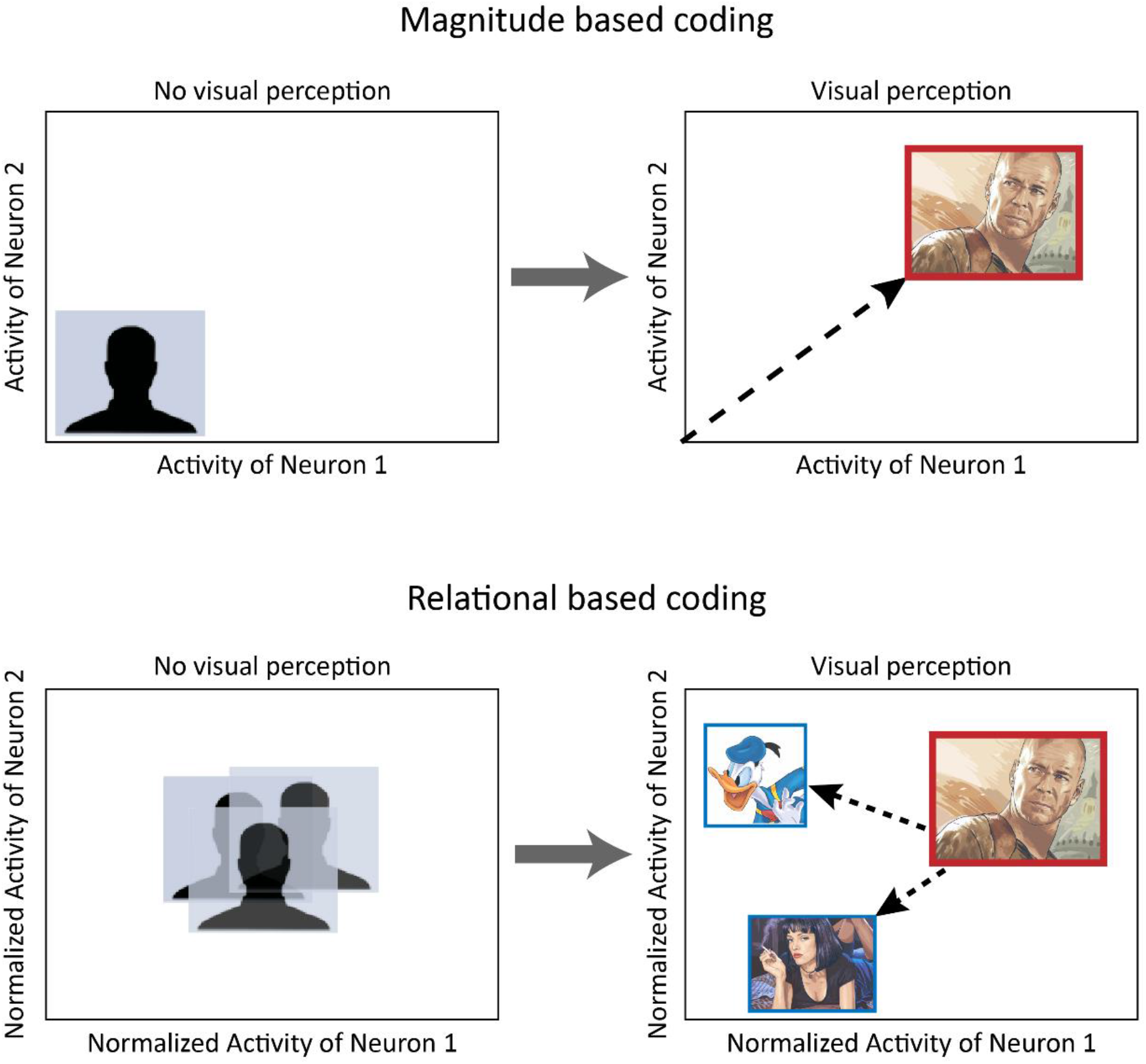
A schematic illustration of two possible visual coding models, depicting the activity levels of two hypothetic visually-responsive neurons. According to magnitude based coding, conscious visual perception derives from an increase in neuronal activity levels. Thus, when the relevant neurons are silent, no visual experience is perceived; when they increase their activity levels, an image, e.g. of Bruce Willis, can be consciously perceived. Relational based coding suggests that an overall amplitude increase in neuronal activity is not enough to generate a unique visual percept. Rather, the distinct visual experience is defined by the correlation-derived distances (or dissimilarities) of the population activation pattern to other activation patterns within the relevant relational geometry-depicted here by dashed arrows in bottom right panel.

This hypothesis extends from previous works that have demonstrated the critical and unique role of pattern-similarities based-coding in visual representations and perception (Edelman et al., 1998; Haxby et al., 2001; Kriegeskorte et al., 2008; Davidesco et al., 2014; Grossman et al., 2019; Kriegeskorte and Diedrichsen, 2019), and have displayed their evolvement across the visual cortex with time (Carlson et al., 2013; Cichy et al., 2014, 2016; Deitch et al., 2021). Here we set out to examine whether these pattern profiles and their similarity distances remain stable across time during extended constant visual stimulation, despite the major reduction in neuronal activation magnitude. Our results also allowed us to test the predictions made in the adversarial collaboration, and hence contribute experimental evidence relevant to two central theories of consciousness in the brain.

## Results

The study is based on intracranial electroencephalographic (iEEG) recordings conducted in 13 patients implanted with 2571 electrode contacts as a part of a clinical diagnostic procedure in the course of treatment for intractable epilepsy (see Table S1 for electrode locations and demographic details of the patients). IEEG recordings were taken while patients performed a simple visual task (see figure 2A and *Methods*), as part of a previous study (Norman et al., 2017; Norman et al., 2019). Twenty-eight images of famous faces and places were presented for 1500ms (14 images from each category), followed by a fixation point on a blank screen for 750ms. Each image exemplar was presented four times in pseudo-random order. The patients were asked to view and memorize the presented images as well as they could, which was verified as highly successful in a subsequent recall session (see Norman et al., 2017; Norman et al., 2019). For analysis purposes, visually-responsive electrode contacts were subdivided into four groups according to anatomical, functional and response latency criteria: early visual electrodes in V1 and V2 (n=32); face-selective electrodes, which showed significantly higher activation in response to faces as compared to places (n=43); content-selective electrodes, that showed a significant preference for a specific subset of exemplars, but not necessarily faces or places only (n=114); and fronto-parietal visually-responding electrodes (n=66); (for further details see *Methods* and Table S1). Figure 2B depicts the location of the iEEG contacts, color-coded according to each group. Note that the face-selective group showed, as expected, some overlap with the content-selective group.

**Figure 2.**
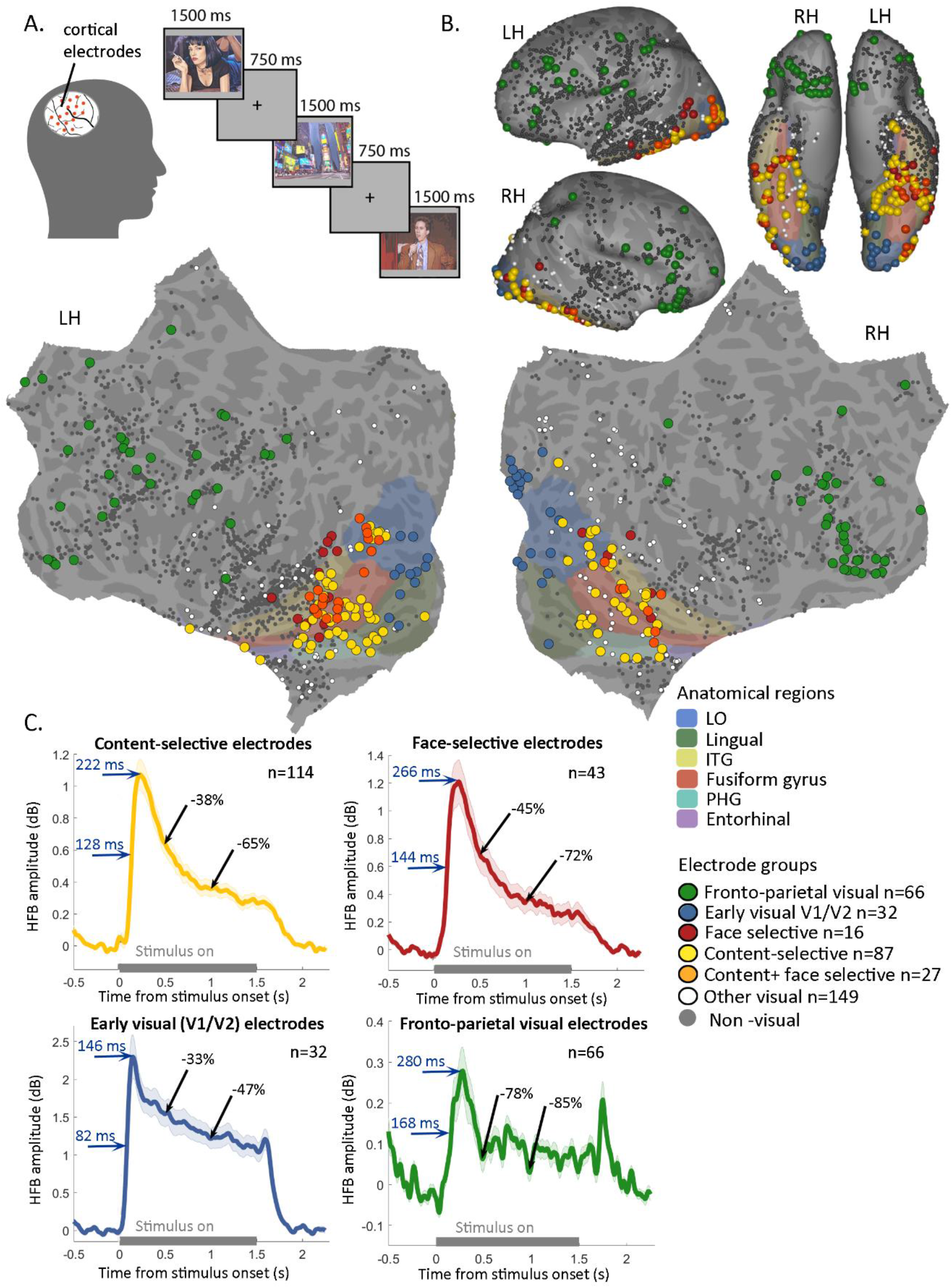
Experimental design, electrode locations, and mean HFB responses to image presentations. **A**. Patients undergoing intracranial recordings viewed images of famous faces and places and were asked to remember them, for a later recall task (Norman et al. 2017, 2019). The stimuli were presented for 1.5 seconds each, with 750ms ISIs. **B**. Multi-patient electrode coverage, shown on inflated (lateral view-upper left; ventral view-upper right) and on flat common cortical surfaces. Electrode locations in each patient were determined based on the post-operative CT and MRI scans, and the cortical surfaces were reconstructed using FreeSurfer and standardized using SUMA, to allow visualization of all the electrodes from the different patients on a shared common cortical template (see *Methods*). Visually responsive electrodes were allocated, based on anatomical and functional criteria to the following subgroups: early visual areas V1 and V2 (blue); face-selective (red); content-selective electrodes in intermediate visual areas (yellow); electrodes that are included both in the face-selective and in the content-selective groups are shown in orange; frontal-parietal electrodes showing visual selectivity (green); and other visual electrodes (white) (see *Methods)*. All remaining contacts, that did not display visual responses, are marked in gray. Colored anatomical labels on the cortical surfaces were derived from surface-based atlases implemented in FreeSurfer (see *Methods*). (Abbreviations: RH—right hemisphere; LH—left hemisphere; LO-lateral occipital cortex; ITG-inferior temporal gyrus; PHG-parahippocampal gyrus.) **C**. Mean HFB (60-160 Hz) responses time-locked to stimuli onset, depicting the average visual response of each electrode set: Content selective electrodes (in yellow), face-selective electrodes (red), early visual electrodes (blue) and fronto-parietal visual electrodes (green). The gray line denotes the stimulus-on duration. Shaded areas represent ±SEM. Blue horizontal arrows point to the half-peak time and peak-time, accordingly, depicting the temporal dynamics of the response build-up. The black arrows denote the percent-signal-decrease of the response, relative to peak amplitude, at times 0.5s and 1s after stimulus onset. Notice the dramatic decrease in the response, i.e. the rapid adaptation effect, that occurs while the stimulus is still presented, across all contact groups.

As a measure of neuronal activity, we have analyzed the broad-band power of high frequencies of the iEEG signal (HFBP: 60-160 Hz), which we and others have demonstrated to be tightly linked to average neuronal firing rates within the recording sites (Mukamel et al., 2005; Nir et al., 2007; Manning et al., 2009). Figure 2C depicts the averaged HFB response magnitude in each contact group, revealing the response latency and the response dynamics elicited by the visual stimulation.

### Response latency across cortical sites

The latency of visual responses across different cortical sites, and specifically the question of a possible delay between high order visual cortex and frontal regions, has been identified as one of the central predictions of the GWT model of conscious perception (Melloni et al., 2021). The wide iEEG coverage in our visual task (see figure 2B) provided a good opportunity to examine this question in the present experimental paradigm. Precise measurements of response latencies for both half-peak and peak response magnitudes (figure 2C, blue arrows) revealed an activation delay between early visual cortex and high order, face-selective contacts of ∼60ms, referring to half-peak times. The half-peak latency of frontal electrodes was of 168ms, but, interestingly, the half-peak latency difference was only ∼20ms between the high-visual face-selective electrodes vs. the fronto-parietal visually responsive electrodes.

### Response magnitude changes during sustained stimulation

As can be seen in figure 2C, all four contact groups displayed a rapid activation onset response triggered by stimulus presentation. Importantly, this rapidly established activation peak only lasted for about 200-300ms and then markedly and significantly declined in all contact groups, despite the persisting, stable visual stimulation. The decline was particularly prominent in the high order content-selective and face-selective groups, decaying to about a third of the peak amplitude within 1000ms. Early visual electrodes showed a higher initial response that declined in a less pronounced manner, while the fronto-parietal visual electrodes exhibited a far weaker and more variable response that decreased prominently, and then, interestingly, exhibited an offset response following stimulus termination.

### Diversity in response patterns and their inter-stimuli distances

Examining the individual contact responses in detail revealed, in accordance with previous observations (e.g. Davidesco et al., 2014; Grossman et al., 2019), a substantial difference in exemplar tuning of different contacts. Complementarily to this diversity of single contact tuning curves, inspecting the multi-contact population patterns of the high-order content-selective electrode group, revealed that each visual exemplar was associated with a distinct profile of a multi-contact pattern, i.e. a unique population vector. This phenomenon is illustrated in Figure 3, which depicts four examples of such multi-contact response profiles and their associated visual images. For the convenience of comparison across these patterns, the electrodes (n=114) are arranged in a descending order of their activation-magnitude in response to the image of Bruce Willis, 300ms after its onset. This serial order of the contacts was maintained in the three other pattern-histograms as well. The electrode colors depict their response magnitude to each specific stimulus, from the strongest response (light blue), to the weakest (black). Two clear aspects can be gleaned from these examples: First, the activation was widely distributed-essentially the entire ensemble of contacts responded at some level to every visual stimulus. Second, the activation levels across contacts were far from uniform, manifested in an approximately four-fold change in activation levels between the highest and lowest responding contacts. Furthermore, as can be seen, each image elicited a unique profile or population activity vector across the set of electrodes, which can readily be appreciated by comparing the different patterns elicited by the four images. Because of the unique association of each pattern with its specific image, one can metaphorically consider such activation patterns as “barcodes”, which mark each visual image with its own unique profile or vector of activations.

**Figure 3.**
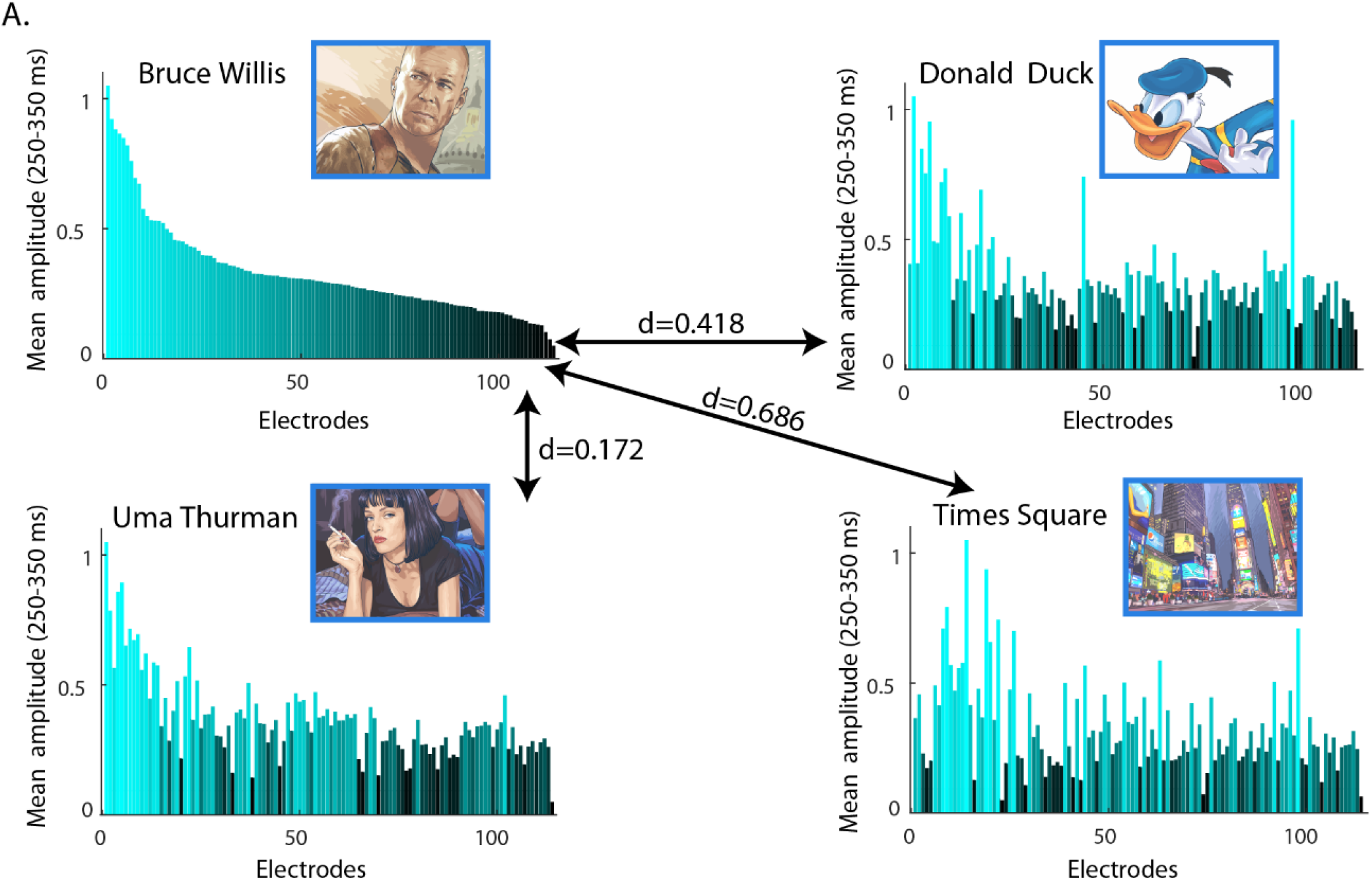
Activity patterns of all visual content-selective electrodes (n=114) to four different stimuli examples. Each histogram depicts the normalized HFB response of all electrodes to the specific stimulus, averaged across time window 250-350ms post stimulus onset. Electrode order is preserved for all four stimuli, and is arranged according to amplitude strength in response to the Bruce Willis image. Electrode color depicts the amplitude level in response to each stimulus, from the strongest response (light blue), to the weakest (black). Distances (d) between the stimuli-evoked patterns are defined as 1 – Pearson’s r, and are depicted by the proportional length of the arrows.

An important point to note is that the similarities between activation-pattern pairs elicited by different image pairs also varied substantially, as can be visually appreciated by comparing the similarity of the Bruce Willis elicited-pattern to the other three activation profiles, depicted in figure 3. The similarities between activity-pattern pairs were quantified in the present study by calculating the Pearson correlation between the two population vectors (see *Methods*). It should be noted that unlike coherence measures, Pearson correlation is sensitive only to the relative activation profiles within the patterns, and not to their absolute magnitude levels. The inverse of the pair-wise pattern similarity was defined as the neuronal *distance* (d) between the pair of stimuli-related patterns, and calculated as 1-Pearson’s r.

### Stability of population activity patterns across time

Relational-based coding, as depicted in figure 1, essentially suggests that each stimulus is embedded in a specific location in the pair-wise pattern-distances vector space. Thus, relational coding is a construct that depends on two components: on the one hand, the unique profile of the activation pattern (the “barcode”) generated by each image, and on the other hand, the unique topology of the relational geometry itself, which can be revealed through the representational dissimilarity matrix - i.e. the matrix of distances between all pair-wise stimuli combinations. Together these factors define, for each stimulus, its relative similarity distances to all other patterns-i.e. its relational code. When examining the neural basis of perceptual stability, an interesting question is whether one of these major components, or both, remain stable during the sustained image presentation durations.

We first examined the stability of the stimulus activation patterns, in the high-order content-selective visual contact group (all results from here on will refer to this set of contacts, until noted differently). Figure 4A depicts the distances matrix, defined as 1-Pearson’s r, between the multi-contact activity patterns across the entire trial duration. Each matrix entry thus indicates the distance between the activity patterns at two time points during the trial (using 100ms averaged time bins), calculated separately per stimulus, and then averaged across all stimuli. In the rest of the text we will term the correlation between activity patterns across time points-*the intertemporal correlation* (see *Methods*). Figure 4B shows the inter-temporal correlation profiles, essentially an inverse of a horizontal cross section line of the distances matrix shown in 4A, at a few example time points. As can be seen, there was a significant increase in the inter-temporal pattern correlations throughout the entire image presentation duration (statistical significance assessed using a 1,000 random-shuffling permutation test, p<0.005, fdr and cluster corrected; significant distances are shown in 100% opacity in figure 4A, and denoted with a yellow line in 4B). The inter-temporal correlations between the early onset-related time windows (200-400ms after stimulus onset) were higher, as compared to later time points in the trial. This transient increase later subsided, revealing stable inter-temporal correlation time-courses at subsequent time points (see the correlation profile at 0.3 seconds as compared to the 1.5 seconds profile in 4B).

**Figure 4.**
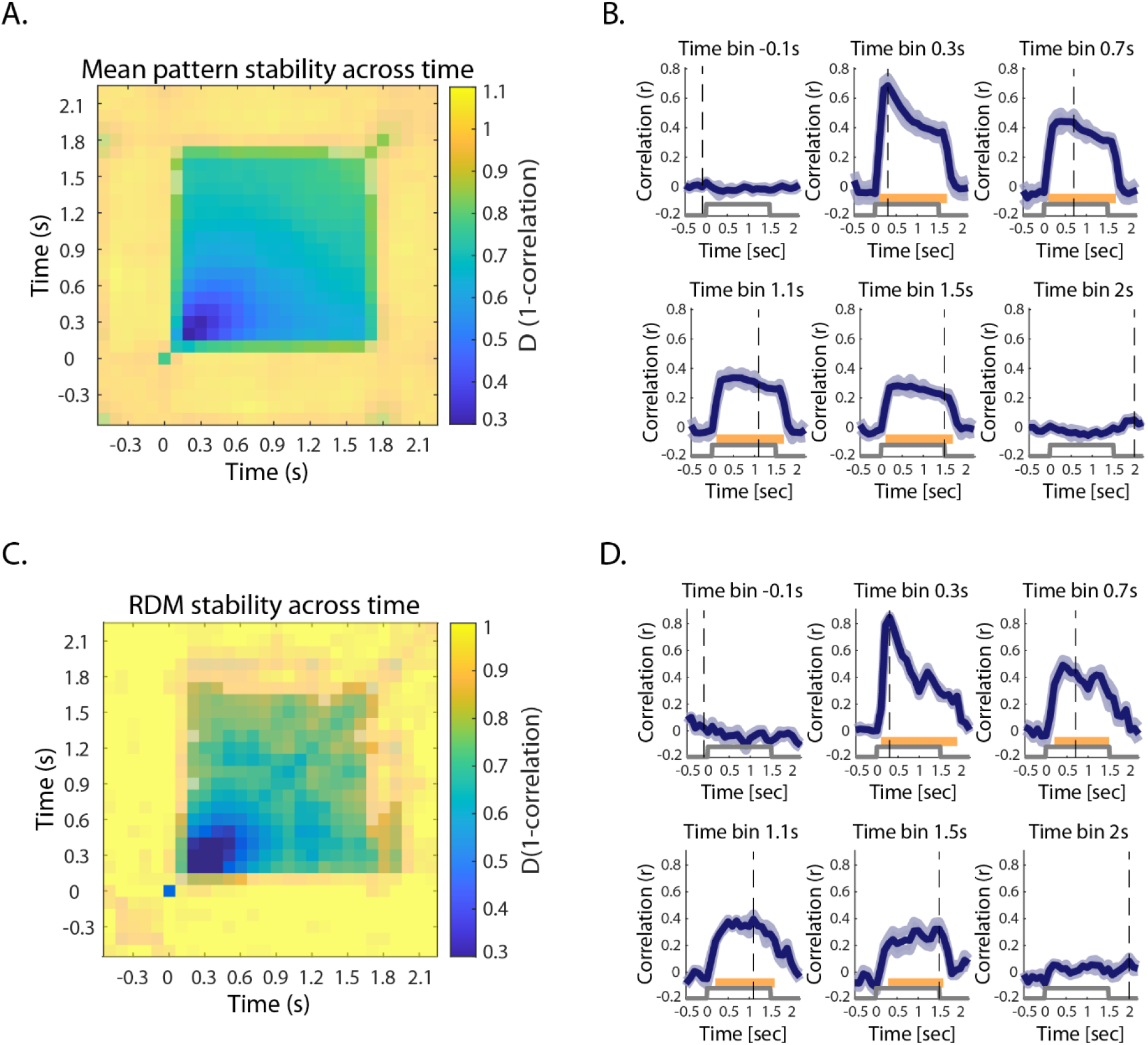
Multi-electrode pattern and RDM stability across time in content selective electrodes. **A**. Mean intertemporal pattern-distances matrix, exhibiting the distances (defined as 1 minus the correlation coefficient) between stimulus-evoked response patterns for all time-bin pairs, using a 100ms time window. Inter-temporal pattern distances were calculated separately for each stimulus (in a leave-1-out manner across the 4 image repetitions), and then averaged across all stimuli. Half-transparent colored regions mark distances that are non-significant, as tested relative to distances emerged from random shuffling, while distance values shown in full opaqueness are statistically significant (1,000 random-shuffling permutations, p<0.005, fdr and cluster corrected), displaying pattern stability across stimulus duration. **B**. Example pattern correlation time-courses, displaying the Pearson-r coefficients between stimuli-related activity patterns at the indicated time bins (−0.1,0.3, 0.7,1.1, 1.5 and 2s relative to stimulus onset) vs. all time bins between -0.5 to 2.2s. Gray step-plots mark stimulus duration time (0 to 1.5 s), and the dashed vertical lines indicate the selected time-bin. Yellow lines indicate statistical significance at p<0.005, fdr and cluster corrected (using 1,000 shuffled permutation test). **C**. Inter-temporal RDM distances matrix, exhibiting the distances between RDMs calculated separately for each time bin, across all time bin pairs. Half-transparent colored regions mark distances that are non-significant, while distance values shown in full opaqueness are statistically significant (1,000 random-shuffling permutations, p<0.005, fdr and cluster corrected), displaying RDM stability across the stimulus duration. **D**. Example RDM correlation time-courses. Gray step-plots mark stimulus duration time, and the dashed vertical lines indicate the selected time-bin. Yellow lines indicate statistical significance at p<0.005, fdr and cluster corrected (1,000 shuffled permutations test).

### Stability of relational geometry

While the stability of individual stimuli-linked patterns has been shown above, it should be emphasized that this does not ensure that the between-stimuli relational distances, and the geometrical similarity-space defined by them, will be stable as well. For example, if all stimuli-induced patterns had identical profiles, then this profile may remain stable-but the relational geometry will be of no information content and hence could be dominated by temporally uncorrelated noise.

As described above, the relational geometry can be revealed by calculating the pair-wise distances matrix of all possible image pairs – generating a Representational Dissimilarity Matrix (RDM) (Edelman, 1998; Haxby et al., 2001; Kriegeskorte et al., 2008). In order to examine the stability of the relational geometry, we first calculated an RDM for each time point. Next, the RDMs were correlated across all time bin pairs, generating an inter-temporal distances matrix displaying the RDM similarity, or stability, across time (see *Methods* for details). Figure 4C shows this intertemporal distances matrix for the relational geometries at different time points, while figure 4D depicts the profiles of the inter-temporal correlation time-courses at selected time points. The overall picture largely resembled the stability of the activity patterns-displaying significant and sustained RDM correlations across the entire duration of the stimuli presentations, with a transient increase in the correlations during the early time windows (statistical significance assessed using a 1,000 shuffling permutation test, p<0.005, fdr and cluster corrected; significant values are shown in 100% opacity in figure 4C, and denoted with yellow lines in 4D).

### Stability of relational coding

Next, we examined the stability of the relational coding of the stimuli, estimated by calculating the distances of a specific image pattern from all other stimuli-elicited patterns in our set, as is illustrated in figure 1 bottom panel and figure 3. If relational coding underlies perceptual content-as we proposed in figure 1, then we would expect it to be maintained for the entire duration of the stimulus presentation-matching the time-course of the perceptual state.

Figure 5A shows the complete profile of the relational code for the Bruce Willis image shown in figure 1, i.e. the distances of all other stimuli-linked patterns from the Bruce-related activity pattern, derived from the content-selective contact group (n=114). As can be seen, these distances were far from uniform-varying over a wide (>three fold) range. This broad range of pattern similarity distances was a common feature in the relational codes of all stimuli, as can be appreciated from figure 5B, which depicts the average, sorted, similarity-distances profiles of all stimuli (see *Methods*). An anticipated category-related effect is also clearly evident in figure 5A and B: distances between stimuli pairs within the same category were significantly smaller than distances across categories (see also the inset in figure 5B, depicting the mean distances across all stimuli pairs for within vs. between categories; p<0.0001, two-sample t-test).

**Figure 5.**
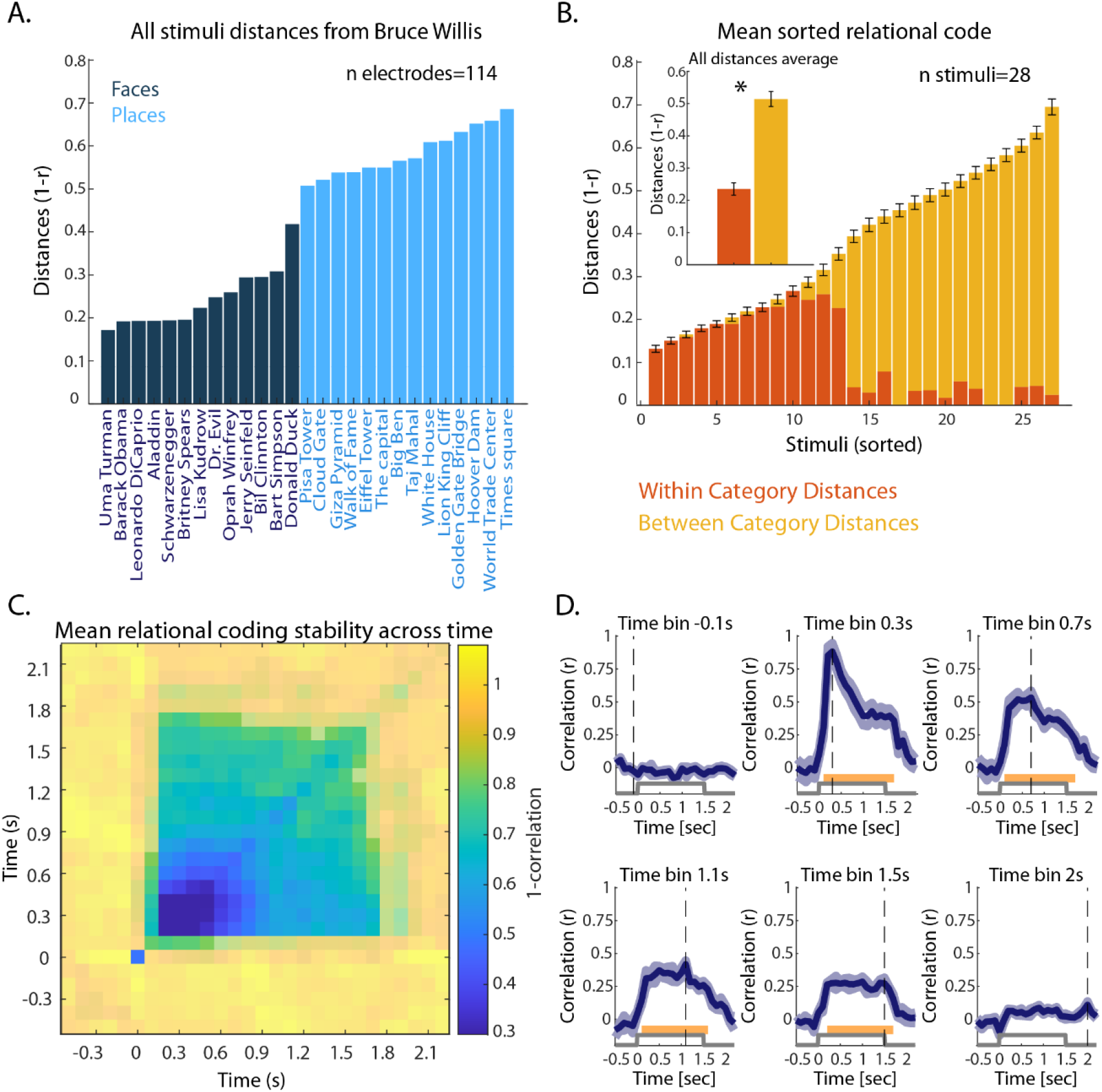
Stability of relational coding across-time, in visual high-order content selective electrodes (n=114). **A**. An example of the relational code of the Bruce Willis image. Histogram bars denote the distances (1-r) of all other stimuli-linked patterns from the Bruce Willis-related multi-electrode activity pattern, at time 0.3s post stimulus onset. Distances are ordered from shortest to longest; distances from face-stimuli patterns are shown in dark blue, and distances from places are in light blue. **B**. Sorted relational code distances averaged across stimuli. For each stimulus, the distances from all other stimuli-linked patterns were calculated and sorted in ascending order, at time 0.3 s (as shown in the example in panel A); these sorted distance vectors were then averaged across all stimuli, to obtain a mean distances vector, illustrating the reliability of this distances-metric across all stimuli. The error bars denote SEM across sorted distances. The proportions of within-category stimuli pairs vs. between category pairs are marked by the relative orange (within category) vs. yellow (between categories) area of each bar, reflecting that within-category stimuli pairs display shorter distances than between-category pairs. This is further emphasized in the inset bar plot, presenting the mean of all within vs. between-category pattern pairs distances (p<0.001, t=-33.97). Error bars denote SEM. **C**. Mean inter-temporal relational coding matrix, exhibiting the distances between the relational code vectors across different time points, averaged across stimuli. Half-transparent colored regions mark distances that are non-significant, as tested relative to distances emerged from random shuffling, while distance values shown in opaque colors are statistically significant (1,000 random-shuffling permutations, p<0.005, fdr and cluster corrected), displaying stability of the relational codes across stimulus duration. **D**. Example intertemporal relational coding correlation time-courses, between 6 selected time-bins and all other time bins. The gray step-functions mark stimulus duration time, and the dashed vertical lines indicate the selected time-bin. Yellow lines indicate statistical significance at p<0.005, fdr and cluster corrected (1,000 shuffled permutation test).

To examine if indeed the relational coding remained stable over the duration of the stimulus presentation, we first calculated the neural distances vector for each individual stimulus, containing the distances between the pattern activity of the specific stimulus to all other stimuli-linked patterns (similar to figure 5A), at each 100ms time bin separately. We then calculated the inter-temporal correlations of these vectors across all time points for each stimulus, resulting in an inter-temporal distances matrix per stimulus. Finally, these matrices were averaged across stimuli in a similar manner to the activity pattern analysis shown in figure 4 (see *Methods*). Figure 5C depicts the inter-temporal matrix of mean relational coding distances, and figure 5D displays a few selected time-point examples of relational-coding inter-temporal correlation time courses. As can be seen, the general dynamics of the relational coding distances across-time were quite similar to the activity patterns and RDM findings, showing a significant and sustained increase in similarity during stimulus duration, with enhanced correlations in the early (200-500ms) response phase. Interestingly, the inter-temporal correlations of the relational code appeared to be slightly more stable compared to the pattern and geometry (RDM) measures in the late phase of the response (see the inter-temporal correlation profiles of time bin 1.5 in figure 5D as compared to figure 4B and D). However, more data will be needed to validate this apparent trend.

### Dynamics and effects of signal to noise levels

What could be the source of the stronger inter-temporal correlations observed during the early phase of the response? Given the higher response magnitude at this early phase, and the possible noise inherent in iEEG recordings, stemming from e.g. muscle and tissue motion artifacts, a likely source contributing to the transiently increased correlations may be a higher signal-to-noise level in the initial, as compared to the later phase of the response. Examining the dynamics of the SNR indeed revealed a major increase in SNR at the initial time windows (see supplementary figure S1A), coinciding with the response amplitude increase, as well as a slight noise decrease. To further examine whether SNR could potentially affect the inter-temporal relational coding correlations measured in our recordings, we compared the inter-temporal correlations derived from time-courses averaged across multiple trials, vs. the noisier single-trial based time course correlations (see *Methods*). This comparison revealed, as expected, that reducing the noise levels (i.e. by averaging across multiple trials) indeed resulted in higher inter-temporal correlation values that were sustained throughout the response period (see supplementary figure S1B and C).

### Pattern stability in high and low order cortical areas

So far, we have examined results from the content-selective contact group, which included the largest number of visually-responsive iEEG contacts in our dataset. In addition to these contacts, we have also obtained a substantial number of recordings from both earlier areas in the visual hierarchy (early visual cortex, n=32) as well as fronto-parietal cortex (fronto-parietal visual contacts, n=66, see figure 2B for the anatomical localization of these contact groups). It was of interest to examine to what extent the stability of the pattern and relational geometries observed in the content-selective group was a general cortical property, or differed across different stages of the cortical hierarchy. Additionally, it should be noted that the content-selective set encompassed a relatively heterogeneous group of visual contacts, including face-selective and place-selective contacts, and contacts that were selective to a mixed set of stimuli from both categories. In order to rule out possible effects due to category-preference differences in this diverse set, we also defined a smaller, more homogenous group of high-order face-selective contacts (n=43, see *Methods*), that partially overlapped with the content-selective group (see figure 2B for their locations).

Figure 6 depicts the inter-temporal distance matrices of the activity patterns, RDMs and relational coding for the face selective, early visual and fronto-parietal contacts (6A, B and C respectively, contact locations shown on inflated cortical surfaces on the left). As can be seen, the face-selective group showed a high resemblance to the content-selective group results, with high intertemporal stability in all three measures (statistical significance assessed by 1,000 random-shuffling permutations, p<0.005, fdr and cluster corrections). Interestingly, it displayed an increase in the stability of the relational coding as compared to the RDM stability across time.

**Figure 6.**
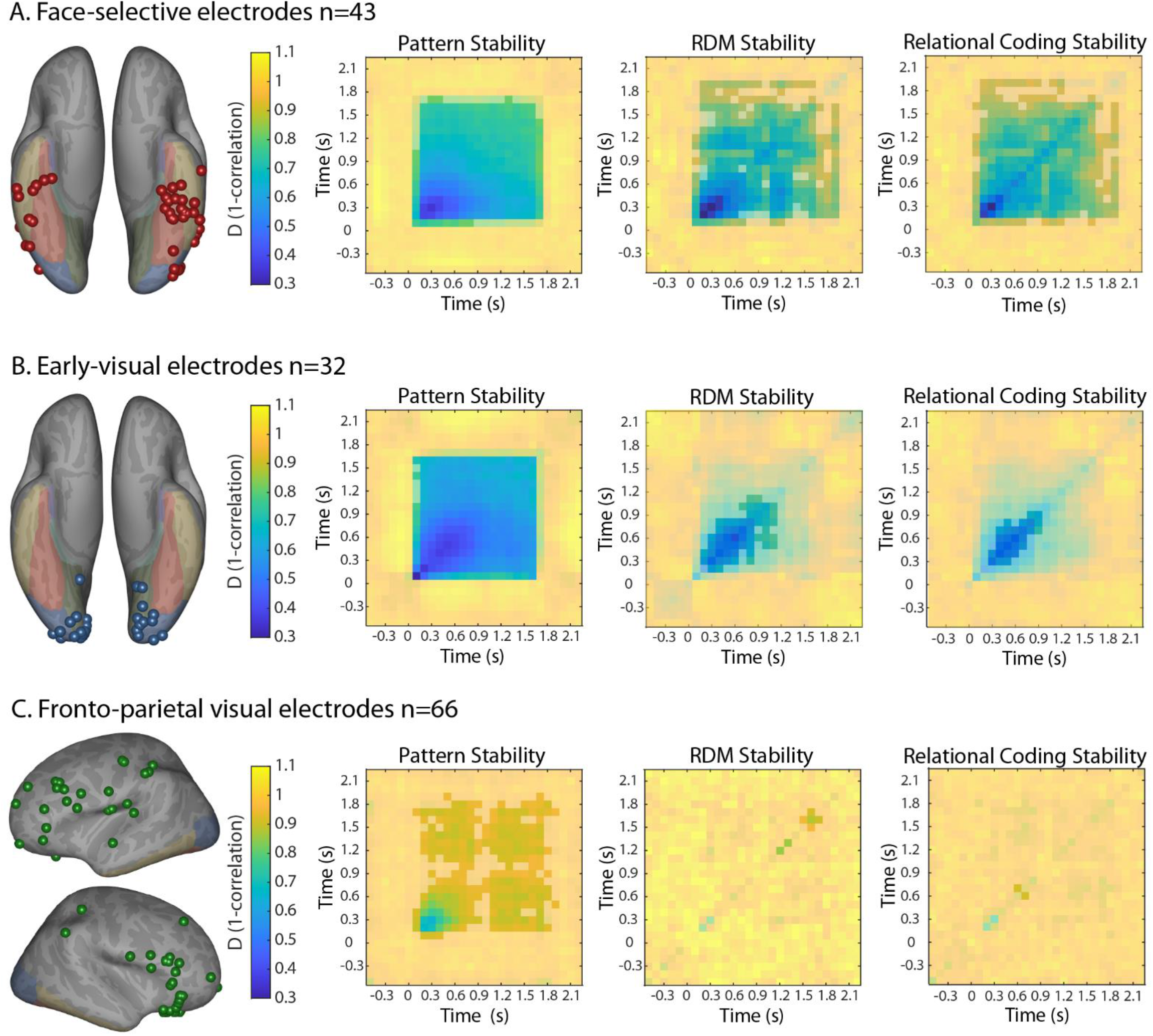
Mean pattern, RDM, and relational coding stability across-time in 3 additional visual functional groups. Electrode locations are presented on inflated cortices in the left-side panels, for face-selective (**A**), early-visual (**B**) and fronto-parietal visual electrodes (**C**). Inter-temporal distances (1-r) of stimuli-evoked patterns, RDMs and relational coding are calculated similarly to figures 4A, 4C and 5C, respectively, for each of the ROIs. Half-transparent colored regions mark distances that are non-significant, while distance values shown in opaque colors are statistically significant (1,000 random-shuffling permutations, p<0.005, fdr and cluster corrected).

By contrast, the early visual group manifested a robust pattern stability effect, but had variable and unstable RDM and relational geometries across-time, displaying significant similarities in the early phase of the stimuli presentations, that were not maintained at later time points. Thus, the relational-geometry representations of the stimuli were not sustained during extended constant visual stimulation in early visual cortex.

The visual fronto-parietal contacts displayed the most prominent differences with essentially no significant inter-temporal correlations of the relational geometry or relational coding, although some weak pattern correlations were observed, mostly around stimulus onset. Various attempts at examining specific subsets of fronto-parietal visual contacts again revealed weak transient patterns at stimulus onset and offset in lateral-frontal contacts, but no significant indication of inter-temporal RDM or relational coding stability (see supplementary figures S2 and S3).

Could the difference between the pattern stability and RDM transiency in early visual cortex be due to less distinct individual pattern profiles-i.e. smaller pattern similarity distances between different stimuli in early cortex, as compared to high order contacts? To examine this possibility, we calculated the across-time averaged distances between activation patterns in early visual contacts and in the high order content selective contacts. The results revealed that the similarity distances in early-visual cortex contacts were significantly smaller than the distances in high-order content-selective contacts (two-tailed paired t-test, t_405_=-10.37, p<0.001, see supplementary figure S4 for the distributions of these distances).

### Stability of pattern decoding

An index of the informational stability that is maintained in the pattern responses across time can also be gained by examining the possibility of inter-temporal decoding, of both the category and the single-item identity of the presented visual stimuli. In fact, inter-temporal category decoding was chosen by the COGITATE adversarial collaboration as one of the decisive measures in contrasting the predictions of the Global Workspace vs. IIT theories (Melloni et al., 2021). In this analysis, a separate classifier was trained for each individual time point, and was then tested across all other time bins, using simple template-matching classifiers for exemplar decoding and k-NN classifiers for category decoding (see *Methods*). The results for these inter-temporal decoders are depicted in figure 7. As can be seen, high order contacts (content and face selective) showed similar results of significant inter-temporal decoding accuracy levels across all training vs testing time bin combinations during the stimulus presentation, though the face-selective contacts showed weaker decoding levels, likely due to the smaller number of contacts in this group. Single-item decoding chance level was 3.57% (1/28), and 50% (1/2) for category decoding, indicated in the figure by the red dashed lines. Further statistical assessment was achieved by using 1,000 shuffled-label permutations, fdr and cluster corrections at p<0.005. Thus, activity patterns in high order visual cortex maintain significantly decodable exemplar-specific and category information across the entire sustained stimuli presentation duration.

**Figure 7.**
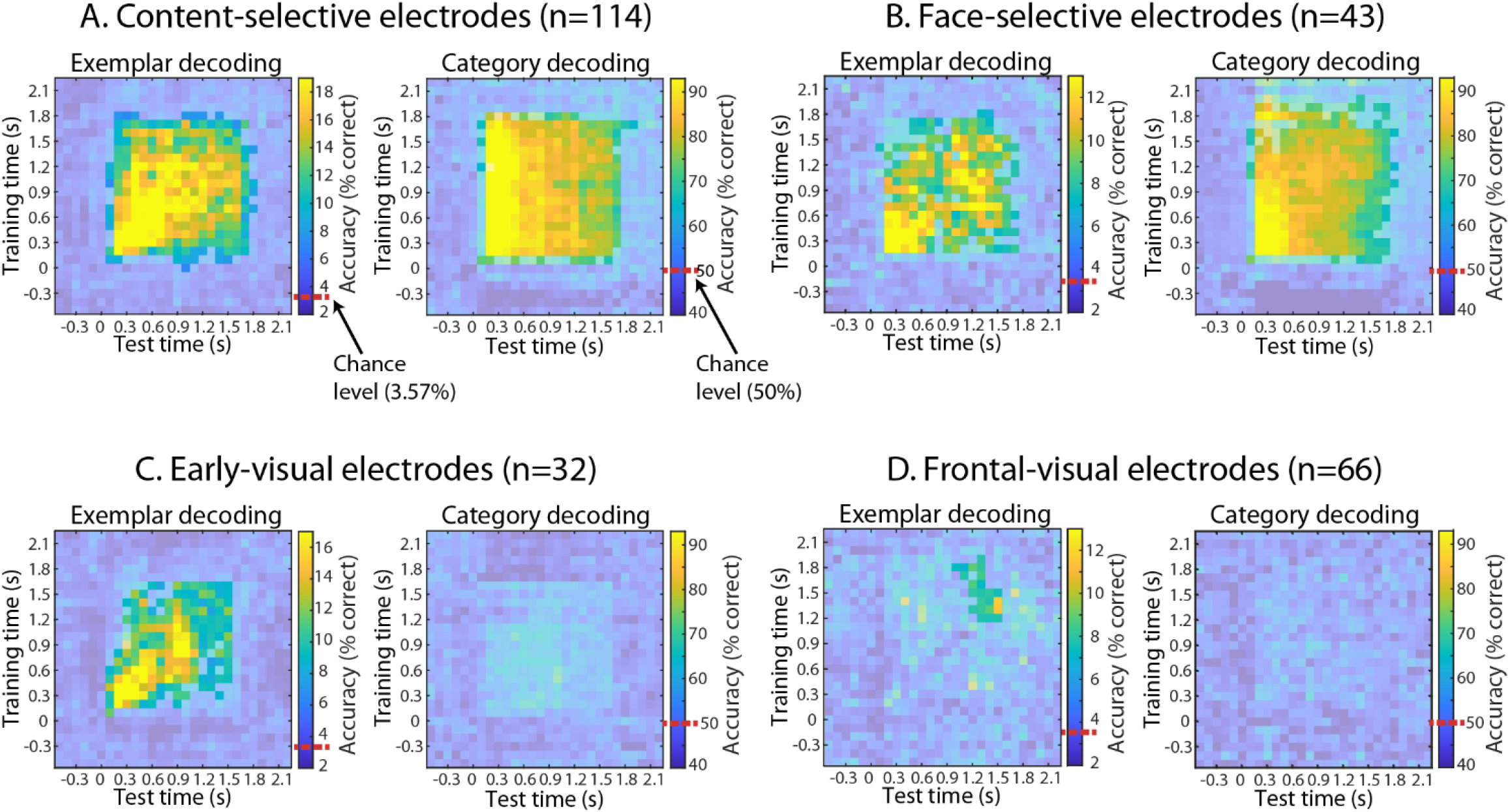
Exemplar and category decoding accuracy across time, in the 4 functional groups: content-selective electrodes (**A**), face-selective electrodes (**B**), early-visual electrodes (**C**) and fronto-parietal visual electrodes (**D**). An individual classifier was trained on the activity patterns of each time bin, and then tested on all other time bins, using simple pattern-matching decoding for the exemplar decoding, and k-NN classifier for the category decoding. Dashed red lines on the color scale bar mark chance level (1/28 = 3.57% for exemplar decoding, and 1/2= 50% for category decoding). Significant accuracy levels were calculated from a shuffling permutation test, comparing the real mean accuracy levels to the distribution of mean accuracy scores from 1,000 shuffled-labels permutations. Statistically significant decoding-accuracy levels are shown in opaque colors (p<0.005 fdr and cluster-based corrections), while insignificant values are shown as half-transparent. See *Methods* for further details.

In order to rule out category-related effects in single-item classification, possibly driven by the different category-preferring contacts in the content-selective group, we additionally trained and tested separate exemplar decoders for each category class individually. These results are presented in figure S5, with both classes showing significant exemplar decoding accuracy above chance level (1/14 stimuli in each category =7.14%) across most of the stimuli duration, though more variable and less stable, compared to the previous exemplar-decoding results. Therefore, inter-temporal exemplar decoding within individual categories is attainable as well, further demonstrating that population activity maintains exemplar-specific information across time.

Inter-temporal decoding results in early visual contacts (figure 7C), interestingly indicated accurate decoding levels only for specific exemplars, with essentially no statistically significant category decoding. Finally, frontal visual electrodes, although showing significant visual responses at the single contact level, failed to show significant inter-temporal decoding either for exemplars or for categories (figure 7D).

## Discussion

### A dissociation between perception and activation magnitude

A dominant outcome of numerous experiments has led to the pervasive conclusion that a strong correlate of the content of perceptual awareness is a significant increase in the activity of the relevant neuronal groups. However, while such high magnitude “ignition” responses (Grill-Spector et al., 2000; Grill-Spector et al., 2004; Fisch et al., 2009; Moutard et al., 2015) may be necessary for the initiation of a perceptual event, the phenomena of rapid visual adaptation challenges the notion of high activation magnitude as a sufficient neuronal correlate of perceptual content. Specifically, in agreement with prior studies (Golan et al., 2016; Gerber et al., 2017) and as shown in figure 2C, during constant visual stimulation, of even a rather short presentation time (<1.5 s) in which no change in perceptual awareness can be discerned, the magnitude of the visual neuronal responses in visual cortex as well as in fronto-parietal regions declines dramatically, dropping to about 70% of the original amplitude in high order visual areas. Thus, it is clear that the magnitude of neuronal activation is unlikely to serve as the neuronal correlate of persistent conscious perception. However, the rapid adaptation phenomenon offers an opportunity to identify such correlate, by searching for aspects of neuronal activity that persist throughout the stable perceptual duration. It is likely that such stable parameters, rather than the magnitude of the responses per se, constitute a more functional neuronal correlate of perceptual awareness across time.

### Stable profiles of activity patterns revealed in high order visual areas

A potential alternative to the notion that the magnitude of neuronal activation is the correlate of perceptual awareness may be found in the profile of the activation patterns. This possibility is compatible with an attractive hypothesis that has been put forward (Edelman, 1998; Edelman et al., 1998; Kriegeskorte and Diedrichsen, 2019; Ebitz and Hayden, 2021; Malach, 2021) proposing that the neuronal mechanism that may underlie conscious perceptual content is not the magnitude of activation, but rather the profiles of neuronal population activity patterns and particularly their unique set of similarity distances to other patterns, defining what we termed here a relational code. Thus, a straightforward prediction of this hypothesis is that the profiles of activation patterns as well as their relational codes, which define the unique location of each stimulus-linked pattern in the representational vector space geometry, will remain unchanged throughout the duration of the perceptual event and will decline rapidly when the image presentation is terminated.

Here we examined this hypothesis in the context of an image viewing task, in which familiar face and place images were presented to the patients for 1.5 seconds (thus long enough for a clear adaptation response, yet still a fairly brief time period). Our results demonstrate that, while activation magnitude rapidly declined within less than 500ms, the similarity of pattern profiles across time (defined through correlation-based distances), the stimuli representational geometry, or RDM, inter-temporal similarity, and the inter-temporal relational code similarity, were all sustained across time for the entire stimulus duration, as is reflected in their significant intertemporal correlation levels. Importantly, this general stability effect was specific to high order content-selective as well as face selective contacts, but not as marked in early visual areas, and not significant in fronto-parietal cortex (see figures 4-6 and supplementary figures S2 and S3). These significant inter-temporal correlations in high-order visual cortex increased rapidly upon stimulus onset and were abolished upon its termination.

Examining the magnitude of inter-temporal correlations revealed a clear and transient increase in the early time points (300-500ms) (Figure 4 and 5). It could be argued that this initial enhancement in correlation values suggests the existence of two different underlying patternresponse profilesan initial “novelty” profile that is then transformed into a different, late onset, pattern profile. However, careful examination of the inter-temporal correlation dynamics at different time points argues against this interpretation. In particular, examining the profile of inter-temporal correlations to the late pattern at 1.5 seconds (figure 4B and D, figure 5D) revealed that it remained stable across the entire duration of the stimulus. Had there been an additional late activation pattern profile, different from the early onset one, we would expect a decline in correlation as we move back in time, from the pattern at the 1.5s time point, to the early time points in the response (i.e. a mirror image of the correlation time-course of the 0.3s pattern). A more plausible explanation of the decline of the early transient peak in the correlation may be a corresponding reduction in signal to noise-which is expected to reduce the measured intertemporal correlations (see figure S1). Examining the SNR over time (figure S1A) showed that it indeed closely followed the change in correlation values. Figure S1B and C demonstrate the clear effect of SNR levels on the inter-temporal correlations of the relational code, by comparing singletrial correlations to across-trials averages. This supports the notion that the underlying pattern profiles remained unchanged along the entire duration of the stimulus.

The inter-temporal correlations of these three pattern-representation measures in high-order visual cortex nicely demonstrate that pattern-based stability is evident at all levels: in the individual stimuli-linked patterns, in the entire stimuli-space geometry defined by the distances between these patterns, and in the specific relational coding of each stimulus within the image vector space.

### Cortical specializations reflected in inter-temporal correlations

We have previously proposed that different cortical regions may manifest distinct relational geometries, endowing them with their unique functional specializations (Malach, 2021). Here we could extend this notion to the temporal domain and examine whether different cortical regions show specialized inter-temporal pattern similarity dynamics over time. Comparing the four visually-responsive contact groups that were defined: early visual, content-selective, face-selective and fronto-parietal, indeed revealed substantial differences across areas. Early visual cortex was characterized by high and persistent inter-temporal activity-pattern correlations, but far weaker and transient RDM and relational coding stability. On the other hand, face-selective and content-selective contacts in high-order visual cortex showed significantly stable pattern and relational geometry inter-temporal correlations across the entire stimulus duration (with the exception of the initial high magnitude correlations). These results are in line with previous studies which combined MEG and fMRI, demonstrating early and transient relational information coding in V1, vs. more sustained RDM correlations in IT (Cichy et al., 2014). Finally, fronto-parietal cortex showed some weak but significant inter-temporal pattern correlations (see figure 6C left panel), but no inter-temporal RDM or relational coding correlations (see figure 6C middle and right panels). However, it should be noted that the visual activations in fronto-parietal cortex were markedly weaker overall, compared to the more posterior regions, so the failure to observe significant inter-temporal correlations may, again, be due to low SNR. We thus cannot rule out the possibility that a larger set of contacts in fronto-parietal cortex may have uncovered significant inter-temporal correlations.

The observed contrast between the stable activity patterns and the unstable relational geometry in early visual cortex regions may appear paradoxical at first, since the representational similarity distances are derived from these patterns. However, such effects can emerge if the patterns associated with different stimuli are not sufficiently distinct from each other-i.e. the similarity distances are on average smaller. In such a case the patterns themselves may show robust stability, but the dissimilarity distances will not be large enough, resulting in RDM and relational coding matrices that are dominated by uncorrelated noise, thus resulting in weak inter-temporal correlations. Directly examining this issue indeed revealed on average smaller pattern dissimilarities in early compared to high order contacts (see supplementary Figure S4). Additional methodological issues in the early visual cortex, e.g. the effects of fixational eye movements or the impact of the retinotopic magnification factor, which are less pronounced in high order visual areas (Leopold and Logothetis, 1998; Golan et al., 2017), may have further influenced these results.

### Stability of relational coding in face-selective neurons

Due to methodological limitations inherent in iEEG, our sampling was relatively limited when confined to a single-category tuned visual electrodes subset, necessitating combining multiple-categories within our main, content-selective, set of contacts. Thus, it could be argued that the relational coding stability we observed in the content-selective contacts was a consequence of combining many different cortical areas and relational geometries within a single population. To examine the validity of this concern we analyzed, in a separate group, the contacts that showed selectivity to a single category-i.e. face-selective contacts. A previous study in our group (Grossman et al., 2019) as well as others (Kriegeskorte et al., 2008) suggested that these contacts comprise a separate, functionally defined, cortical geometry.

While the results from this face-selective contact group were noisier, likely due to the substantial reduction in the number of contacts (114 in the content-selective group vs. 43 in the face-selective group), they were essentially the same, exhibiting sustained inter-temporal correlations in pattern activity, RDM and relational coding stability.

### Inter-temporal decoding and its relevance to dominant theories of conscious experience

Recently, in an attempt to experimentally examine the predictions of two central consciousness theories – Global Workspace Theory and Integrated Information Theory, an “adversarial” collaboration proposed to examine inter-temporal visual information stability as an experimental test of these theories (Melloni et al., 2021). Specifically, GWT predicts that fronto-parietal regions should show unique inter-temporal stimulus category decodability dynamics, with high decoding levels around stimulus onset and offset, that decline to zero between these times. By contrast, IIT proposes that category decodability will be sustained across the entire length of the stimulus duration and be localized to posterior visual cortex.

Our paradigm provided an opportunity to experimentally test these predictions. We found that by contrast to the GWT prediction, the visual fronto-parietal contact group did not exhibit any above-chance inter-temporal decoding levels for stimulus category. However, it is important to note here that decoding is a measure that combines both a relational code (how similar is the trained pattern to the tested one) and noise level (how variable is the activation). Given that the frontal responses in our study were markedly weaker than the posterior activations, we cannot rule out at this stage that some significant inter-temporal decoding may be feasible in fronto-parietal contacts if a larger number of recording contacts will be available.

It should be noted here that our latency measurements were also incompatible with the GWT prediction of around 100ms activation lag between posterior visual areas and the fronto-parietal ignition (Melloni et al., 2021). Our results reveal a latency lag within the visual system proper-i.e. between early and high order face-selective regions, but only a minimal latency lag (∼20ms) between high order face-selective cortex and the fronto-parietal response, as can be seen in figure 2C, which is also compatible with our previous results from large scale intra-cranial recordings (Noy et al., 2015).

As to IIT predictions, our results appear to be compatible at the qualitative level with their prediction of significant and persistent inter-temporal decoding in the visual cortex. However, it should be emphasized that such persistent effects are predicted also by non IIT hypothesis (e.g. Malach, 2021).

It should be noted that because pattern decoding is partially based on relational coding, our decoding results provide further support to the suggestion that different cortical regions manifest different relational geometries, reflected, for example, in the striking difference we found in category decoding between high and low order visual areas (see figure 7). This effect can be readily explained by the likely specialization of early visual cortex neurons in local visual features (Lerner et al., 2001), which were common across the face and place images in our set.

### Possible roles for high neuronal activations

The present findings raise the question of the role of the early, transiently high, neuronal activation levels, which are a common observation in visual responses. These transient bursts, termed “ignitions”, are a consistent signature of crossing the conscious perceptual threshold (Grill-Spector et al., 2004; Quiroga et al., 2008; Fisch et al., 2009; Moutard et al., 2015). At present, the function of these high activity bursts remains an open question-however, there are a number of attractive possibilities, which in fact may complement each other. It was recently proposed (Malach, 2021) that the high firing rates associated with the initial onset of conscious perception may implement a binding mechanism that integrates the individually active neurons into a joint population with a coherent relational geometry. It should be noted that since neuronal information is likely coded in the dynamically changing firing frequencies-e.g. inter-spike intervals, high activation rates have the advantage of rapid binding and information transfer. However, as discussed here, this raises the issue of stability-i.e. how can the neuronal binding be maintained if the binding mechanism itself is so transient. One possibility is that following the rapid formation of the relational geometry, its maintenance may require only low-levels of activity (Barak and Tsodyks, 2014). Additional potential functions of the initial high firing could be its rapid spread to down-stream cortical areas, its registration in working memory, as well as in long term memory and in rapid motor actions. Clearly, future studies are needed to resolve all these fundamental questions.

## Conclusions

In summary, our study points to relational coding as a plausible mechanism that enables neuronal representations to remain stable across prolonged time periods. This may account for the sustained conscious percept elicited by constant external stimulation, despite the massive changes in the magnitude of activation. This relational coding mechanism is based on the profile of multi-neuronal activation patterns and the unique geometry defined by their pairwise distances, and is expressed most reliably in high order, content-selective, visual cortex.

## Acknowledgments

We are grateful to the patients for their kind cooperation Funding: Supported by U.S.-Israel Binational Foundation grant 2017015 (R.M. and A.D.M.) and a CIFAR Tanenbaum Fellowship (R.M.).

## Methods

### Participants

Intracranial recordings were obtained from 13 patients (10 females, mean age 34.7 ±9.6), monitored for pre-surgical evaluation of epileptic foci due to pharmacologically resistant epilepsy, at the North Shore University Hospital in NY. As part of the clinical assessment, all patients were implanted with subdural or depth electrodes (see Table S1 for individual demographic and electrode coverage details). The study was conducted in accordance with the latest version of the Declaration of Helsinki, and all patients provided a fully informed consent to participate, including consent to publish the results, according to the US National Institute of Health guidelines, monitored by the institutional review board at the Feinstein Institute for Medical Research. No epileptic clinical seizures occurred during the experiment.

### Experimental task and stimuli

The participants viewed images of famous faces and places, as part of a longer study that included a later phase of free recall of the images that were presented (Norman et al., 2017; Norman et al., 2019). The experiment was divided into two runs. Each run began with a 200 seconds resting-state period with eyes closed (the first two patients performed the resting-state task on a different day). Immediately afterwards, 14 different images of well-known faces and places were presented to the participants (7 pictures from each category). In total, 28 different naturalistic and colorful stimuli were included in the experiment, 14 in each run (see figures 1,2A and 3 for examples of these stimuli). Each image was presented for 1500ms, with 750ms inter-stimulus intervals (ISIs) between them, during which a fixation cross was presented (see figure 2A for task depiction). Single items were repeated four times each, in a pseudo-random order such that no image was repeated twice consecutively. The stimuli were presented on a LCD screen, using Presentation software (Version 0.70, www.neurobs.com), at a ∼60 cm viewing distance (image size was 16.5° × 12.7). The participants were asked to carefully view these images and try to remember them as well as possible, including details regarding unique colors, facial expressions, perspective, lightning, etc. The free recall task took place after the image viewing phase of each run (see Norman et al., 2017; Norman et al., 2019).

### Electrodes implant and data acquisition

All recordings were conducted at the Northshore University Hospital, Manhasset, NY, USA, at the patients’ quiet bedside. Electrodes were either subdural grids or strips, places directly on the cortical surface, or depth electrodes (Ad-Tech, Racine, WI, Integra, Plainsboro, NJ, and PMT Corporation, Chanhassen, MN). Subdural electrodes were either 1 or 3 mm in diameter, with inter-contact spacing of either 4 or 10 mm. In depth electrodes were 2 mm platinum cylinders, with 4.4 mm inter-contact spacing, and a diameter of 0.8 mm. The intracranial signals were referenced to a vertex screw or to a subdermal electrode, and were electronically filtered between 0.1 and 200 Hz, and then sampled at a rate of 500 or 512 Hz. The data was stored for offline analysis using XLTEK EMU128FS or NeuroLink IP 256 systems (Natus Medical Inc., San Carlos, CA). Stimulus-triggered electrical pulses were sent upon stimuli onsets and recorded along with the iEEG data for precise alignment of task protocol to neural activity.

### Electrode anatomical localization

Before the electrode implantation, the patients underwent a T1 weighted 1 mm isometric anatomical MRI scan using a 3T Signa HDx scanner (GE Healthcare, Chicago, Illinois). Following the implant, a computed tomography (CT) and a T1-weighted anatomical MRI scan on a 1.5-T Signa Excite scanner (GE Healthcare) were acquired, in order to enable precise localizations of each electrode. All scans were skull-stripped using FSL’s BET algorithm, and the post-implant CT was first co-registered to the post-implantation MRI scan, and then to the pre-implantation MRI anatomical scan, using rigid affine transformation, as implemented in FSL’s Flirt (Jenkinson and Smith, 2001; Jenkinson et al., 2002; Smith, 2002). Concatenation of these two co-registrations allowed visualization of the post-implant CT scan on top of the preoperative MRI scan, while minimizing potential errors due to surgery and implantation possible brain shifts. Individual electrodes were next identified by manual inspection of the co-registered CT and post-implant MRI, and marked in each patient’s preoperative MRI native space, using the BioImage Suite (Papademetris et al., 2006). Next, electrode projection onto the cortical surface was performed as previously reported (Golan et al., 2016; Norman et al., 2019): First, individual patients’ cortical surfaces were segmented and reconstructed from the pre-implant structural MRI, using FreeSurfer 6.0 (Fischl, 2012). Then, each electrode was allocated to the nearest vertex on the individual’s cortical surface. Contacts that were farther than 8mm from the cortical surface were excluded from further analyses. In order to project all electrodes from all patients onto a single template cortical surface, while maintaining their specific relations to individual gyri and sulci, the 3-D cortical mesh of each individual was resampled and standardized using SUMA (Argall et al., 2006), allowing visualization of all electrodes on a single common cortical template (FreeSurfer’s FSaverage). Finally, colored labels on the cortical surface, as shown in figure 2B, were derived from different surface-based anatomical atlases available in FreeSurfer (Desikan et al., 2006; Fischl et al., 2008; Destrieux et al., 2010), including a probabilistic visual-retionotopic atlas (Wang et al., 2015).

### iEEG data preprocessing and HFB estimation

All data analysis was performed in MATLAB (MathWorks), using EEGLAB (Delorme and Makeig, 2004), Chronux (Bokil et al., 2010), DRtoolbox (https://lvdmaaten.github.io/drtoolbox/), MES toolbox (Hentschke and Stuttgen, 2011), and in-house developed code. Raw iEEG time-courses were inspected manually and statistically to detect noisy/corrupted channels, which were then excluded from further analysis. Signals that were recorded at a sampling rate of 512 Hz were downsampled to 500 Hz, for consistency. The 60 Hz power line interference, as well as its harmonics, were removed using zero-lag linear-phase Hamming-windowed FIR band-stop filters. Next, all electrodes were re-referenced to a robust common average (excluding the corrupted channels).

The high-frequency broadband (HFB) signal was defined as the mean normalized power of frequencies between 60-160 Hz (High-gamma), the range of frequencies that is commonly used as the electrophysiological marker of neural population activity (Mukamel et al., 2005; Nir et al., 2007; Manning et al., 2009; Norman et al., 2017; Parvizi and Kastner, 2018; Norman et al., 2019). HFB power was computed by filtering the signal into 20 Hz bands between 60-160 Hz, employing zero-lag linear-phase Hamming window FIR filters, using EEGLAB. Next, the momentary amplitude in each sub-range was calculated as the absolute value of the filtered signal’s Hilbert transform (Fisch et al., 2009; Noy et al., 2015; Grossman et al., 2019; Norman et al., 2019). Since the 1/f profile of the signal’s power spectrum results in larger values of lower frequencies, we normalized each sub-range by dividing it with its mean value. Finally, we averaged the normalized values across all sub-ranges. The resulting HFB data was inspected for transient electrical artifacts, defined as signals above 5 SD in the common average signal (the averaged time-series across all electrodes). Time windows of 200ms around these peaks were removed from further analyses.

The HFB data was then epoched relative to stimulus onset (−550 to 2250 post stimulus onset). In order to normalize the data to account for differences in overall HFB amplitude levels in different electrodes, event-related HFB responses to visual stimulation were normalized relative to a baseline period of -400 to -100 ms prior to stimuli onset, by dividing each time point in the epoched data by the mean baseline amplitude. Finally, since the HFB amplitude tends to follow a log-normal distribution, as do many other measures of population firing rate, the HFB values were log-transformed by 10*log10 (Norman et al., 2017; Norman et al., 2019).

### Definition and grouping of visually responsive contacts

Visually-responsive contacts were identified by comparing the post-stimulus HFB response of each contact (averaged across the time window of 100 to 500ms post stimulus onset) to its pre-stimulus baseline (averaged from -400 to -100ms prior to stimulus onset), using a two-tailed Wilcoxon signed-rank test. All contact p-values (from all patients) were pooled together to control for the false discovery rate (FDR) (Benjamini and Yekutieli, 2001). Visually responsive contacts were defined as those that displayed a significant HFB response (p_fdr_<0.05). Next, we grouped together visually responsive contacts based on their anatomical, functional, and response latency features, to the following subsets: early visual (V1/V2), face-selective, content-selective, and fronto-parietal visual contacts.

In order to calculate the response latency of each visually responsive contact, we compared the HFB amplitude in each post-stimulus time point to the pre-stimulus baseline, using a paired t-test. Response latency was defined as the first time point in which the HFB amplitude was significantly higher than baseline (p<0.05), and remained significant for the next 50ms at least (Foxe and Simpson, 2002). Previous single-unit studies in monkeys reported a maximal response latency of ∼180ms in early visual regions V1 and V2 (Raiguel et al., 1989). Thus, we defined early visual contacts (n=32, marked in blue in figure 2B) as the visually-responsive contacts that showed a response latency of 180ms or less, and were located in Brodmann areas 17 and 18 (V1 and V2), as based on the Brodmann atlas implemented in FreeSurfer (Fischl et al., 2008).

To define face-selective contacts, the mean HFB responses across the 100-500ms time window post-stimulus onset, were compared between faces and places using a Wilcoxon rank sum test. Contacts that showed significantly higher activation in response to faces than places (p_fdr_<0.05), and were anatomically located beyond early visual regions, excluding frontal cortex, were defined as face-selective contacts (n=43, denoted in red and orange in figure 2B).

An additional set of visual contacts was termed content-selective contacts, and included contacts that showed preferential responses to specific stimuli exemplars as compared to others; importantly, these were not necessarily only faces or places (based on Norman et al. 2019). In order to define these contacts, we inspected the visually-responsive contacts located in 6 anatomical regions across the ventral visual hierarchy, based on the Desikan Killiany atlas (Desikan et al., 2006), including the lateral occipital cortex (LO), inferior temporal gyrus (ITG), lingual gyrus, parahippocampal gyrus (PHG), fusiform gyrus, and entorhinal gyrus, excluding the early-visual contacts that were already defined. Visually responsive contacts that were located in these ROIs, and showed significant content-selectivity in their responses, defined as a difference of at least 3.5 SD between the top 10 preferred images and bottom 10 images, were included in the content-selective contact set (n=114, shown in yellow and orange in figure 2B). Notice 27 contacts fitted the criteria for both the face-selective group and the content-selective groups, depicted in orange in figure 2B.

Finally, in order to examine the activity patterns in prefrontal and parietal regions, we defined a fronto-parietal anatomical ROI based on the Desikan Killiany atlas labels, that included the superior frontal gyrus, rostral middle frontal gyrus, pars orbitalis, pars triangularis, pars opercularis, precentral gyrus, postcentral gyrus, supramarginal gyrus, orbital frontal gyrus, and the anterior cingulate. All visually-responsive contacts that were located within these regions were included in the fronto-parietal visual group (n=66, shown in green in figure 2B). For additional supplementary analyses, we also subdivided this fronto-parietal group to more spatially restricted visually-selective contact sets: an orbitofrontal cortex (OFC) contacts group (encompassing the lateral and medial orbitofrontal cortex; n=20, see figure S2) and a lateral prefrontal group (including the superior frontal gyrus, middle frontal gyrus, precentral gyrus, pars orbitalis, pars triangularis and pars opercularis; n=35, see figure S3).

Additional visually-responsive contacts that did not fall under one of the above visually-selective subgroups were labeled “other visually-responsive contacts”, and are shown in white in figure 2B; non-visual contacts are marked in gray.

### Population activity patterns’ inter-temporal stability

In order to quantify the stability of the stimuli-evoked population activity patterns across time, we calculated the correlation based distances of these patterns between all time points. First, we split the HFB data into epochs relative to stimuli onset, including the 550ms pre-stimulus fixation, 1500ms stimulus presentation duration, and the 750ms post-stimulus fixation period (resulting in 2800ms long epochs). Each stimulus (n=28) was presented 4 times, thus we had 112 such epochs in total per contact (28 stimuli*4 repetitions). Next, we downsampled the data by averaging it across time in 100ms non-overlapping time windows, hence resulting in 28 time bins per trial (or epoch). Then, for each contact subset (see “*Definition and grouping of visually responsive contacts”* above), we arranged the data into a 3D matrix per each stimulus, resulting in 28 matrices depicting the activity patterns, of the following form: ***A*** _*n contacts × 28 time bins × 4 repetitions*_. Examples of such activity patterns (for the content-selective contact group), for time bin 0.3s post-stimulus onset, are shown for four different stimuli in figure 3 (activity patterns were averaged across the 4 stimulus repetitions in this figure).

For every **A**_i_ matrix (i=1,2… 28 stimuli), we correlated the contact activity pattern vectors between all time-bin combinations, so that for each specific time bin we obtained the Pearson’s r correlation values between the contact patterns of all other time bins. In order to avoid auto-correlations and possible noise artifacts, the correlations were calculated in a leave-1-out procedure across the 4 stimulus repetitions, in the following manner: The correlation values for each time-bin pair were calculated 4 times, so that in every iteration we extracted one trial and averaged the pattern activity across the remaining 3 trials, and then correlated between the averaged pattern and the single-trial pattern. We then proceeded to calculate the average correlation value across these 4 iterations. Distances were defined as 1-Pearson’s r. Thus, we obtained 28 inter-temporal distance matrices (for each of the 28 stimuli), of the following form: ***D***_*28 time bins × 28 time bins*_. Finally, we averaged across the individual stimuli **D** matrices, resulting in a single averaged inter-temporal distances matrix, shown in figure 4A (and in figure 6-denoted as the “Pattern Stability” matrices).

In order to assess the statistical significance of the inter-temporal pattern correlation/distance values, we shuffled the contact labels in the original data 1,000 times, and then proceeded to recompute the correlations in the same manner as described above, resulting in a shuffled 1,000 r-values distribution for each i^th^ time bin vs j^th^ time bin entry. P-values were calculated as the proportion of correlation-based distance (1-r) values derived from the shuffled permutations that were smaller than the original distance value derived from the data (smaller d values indicate higher similarity). An FDR correction (Benjamini and Yekutieli, 2001) was then applied on the resultant 784 p-values (28×28 time-bin pairs), to control for a 0.5% false discovery rate across all time-bin comparisons. In addition, we assessed the maximal cluster size in each shuffled iteration (using a cluster-defining threshold of p=0.05). FWE-corrected p-values were computed as the proportion of random clusters larger than or equal to the clusters observed in the actual data (Maris and Oostenveld, 2007; Groppe et al., 2011; Oostenveld et al., 2011). The distance values that survived the combined FDR and cluster-based corrections were marked as significant in the inter-temporal pattern stability matrices.

### RDM computation and across-time stability analysis

In order to compute the stimuli pair-wise distances matrix, i.e. the representational dissimilarity matrix (RDM), we first arranged the HFB epoched data, for each contact group separately, in neuronal stimulus representation matrices **B**, in the form of ***B*** _*n contacts × 28 stimuli × 4 repetitions per stimulus*_, with a separate **B**_**i**_ (1=1,2…28) matrix for each time bin. The pair-wise distance between the neural response patterns of two exemplars was calculated as 1-Pearson’s r, depicting the correlation between the activity pattern vectors of two stimuli. Distances were calculated between all possible stimuli pairs, separately for each **B**_**i**_ matrix. The correlations were calculated in a leave-1-out manner to avoid autocorrelations and artifacts, as explained in the previous section (*Population activity patterns’ inter-temporal stability)*, iterating across the 4 possible combinations of a single trial vs. 3-trials averaged activity vector, and deriving four correlation values for each stimuli pair, that were saved in separate RDMs. Thus, we obtained a total of 112 RDMs depicting the distances between all possible stimuli pairs, each in the form of ***R***_*28 stimuli × 28 stimuli*_, one for each of the four leave-1-out iterations, and for each time bin separately (n=28). In order to determine the stability of the RDMs across-time, as an indication to the stability of the stimuli representational geometry, we next examined the inter-temporal correlations of these RDMs. In that aim, we first “unrolled” the top and bottom triangular halves of each RDM (including the main diagonal) into vectors, and averaged across them. This resulted in a vector of 406 distance values (*N*∗ (*N*− 1)/2 +*N*), depicting the distances between all stimuli pairs, including the distances between different presentations of the same stimulus (N=28, denoting the number of stimuli). These pair-wise distance vectors were obtained for each time bin, and for each of the four leave-1-out iterations. Next, we correlated these distance vectors between all possible time-bin pairs, again in a leave-1-out-manner, resulting in a 28 time bins × 28 time bins × 4 leave-1-out iterations matrix, which we then averaged across the 3^rd^ dimension. This resulted in a 28 × 28 distances matrix, depicting the inter-temporal stability of the RDM, or stimuli representational geometric space (as shown in figure 4C, and in figure 6- the matrices termed “RDM stability”). Statistical significance was assessed through 1,000 shuffled permutations, using FDR and cluster-based corrections, in the same manner as described above in “*Population activity patterns’ intertemporal stability”*.

### Inter-temporal stability of relational coding

The relational code of a stimulus was defined as the vector of pattern distances from that specific stimulus to all other stimuli. In order to examine the across-time stability of these relational codes, we first extracted RDMs for each time bin separately, as described in the section above (see *RDM computation and across-time stability analysis)*. Every row in these RDMs consists of a distances vector, denoted *d*_*i*_ (i=1,2…28 stimuli) that holds the relational code for a specific stimulus-i.e. the distances between one stimulus and all other stimuli, for a specific time bin. Next, we calculated the across-time correlations for every *d*_*i*_ vector separately, in a leave-1-out manner, as described in the two sections above. This resulted in 3D distance matrices of size *28* _***time bins***_ × *28* _***time bins***_× *4* _***leave-1-out combinations***_ which were averaged across the 3^rd^ dimension, resulting in a *28* _***time bins***_ × *28* _***time bins***_ mean distances matrix for each individual stimulus. Lastly, we averaged across the 28 individual-stimulus distances matrices, resulting in the final grand average *28* _***time bins***_ × *28* _***time bins***_ distances matrix, that reflected the average stability of the relational coding across trial duration (e.g. in figure 5C). This was done separately for every contact group. Statistical significance was assessed through 1,000 shuffled permutations, using FDR and cluster-based corrections, in the same manner as described in the two sections above.

### Signal-to-noise analyses

Signal to noise ratio (SNR) was calculated as the average HFB signal across all trials and stimuli, divided by the standard deviation computed across repetitions of the same stimulus, and averaged across all stimuli. Examining the SNR time-course revealed that earlier time points in the trial, immediately following stimulus onset, displayed higher SNR levels. In order to explore how higher SNR levels can potentially contribute to the increased correlation values we obtained in these time points, we rerun the stimulus relational coding analyses pipeline twice, as described above, but using different SNR levels of the data. This was done by running the analysis once with averaging across individual repetitions of the same stimulus before calculating the correlations (in the leave-1-out manner as described above for the RDM computation, and then by averaging across pairs of RDMs before calculating their inter-temporal correlations); and once without averaging-i.e. calculating correlations between single trials, and only then averaging across the correlation values derived from the different single-trial combinations. Since averaging across trials increases the SNR, in this manner we could visualize the effects of higher SNR levels, which led to higher correlation levels in the relational coding stability analysis.

### Single-exemplar and category decoding across-time

To test whether information regarding the identity of single exemplars, as well as the category of the stimulus, is sustained across time, we trained and tested inter-temporal exemplar and category decoders, separately for each contact group that we defined. First, we applied pattern dimensionality reduction (PCA) to the data, in order to reduce feature number before training the decoders (Van Der Maaten et al., 2009; Norman et al., 2019). This was done by averaging the epoched HFB data across the trial durations, and then across individual stimuli repetitions, constructing a mean feature matrix of the visual responses to all stimuli *(28*_*exemplars*_ *× n* _*contacts*_), and then applying the PCA algorithm on this matrix. In order to determine the optimal number of PCs to retain, we estimated the true dimensionality of the data (i.e. the intrinsic dimension) using a Maximum Likelihood Estimation (MLE) algorithm (Levina and Bickel, 2005). We proceeded to maintain the first 12 PCs, that accounted for 89.2% of the variance in the data for the content-selective contact group. Next, we applied the linear transformation obtained from the PCA to the original data (using an out-of-sample extension of the PCA), resulting in a 12-dimensional linear space, which was arranged in a grand four-dimensional matrix *G* of the form: *12*_*PCs*_*×28*_*time bins*_*×28*_*exemplars*_*×4*_*repetitions per exemplar*_. Thus, entry *G*_*i,j,k,l*_ refers to the response level of the i^th^ PC, at the j^th^ time bin (100ms bins), for the k^th^ exemplar and the l^th^ repetition trial of that exemplar.

In order to train the inter-temporal single-exemplar decoders, we applied a simple template matching decoding technique, in the following manner: We looped over all individual time bins in two nested loops, in order to train and test every possible combination of training a decoder on time bin *m*, and testing it on time bin *n*, for a total of 28^2^=784 time-bin combinations. For each unique training vs. testing time-bin combination, we ran 1,000 decoding iterations. In each iteration, we randomly chose a single trial for each exemplar, extracted the response it elicited at time bin *n* from all PCs, and assigned it to a test pattern 2D matrix *T (28*_*exemplars*_*×12*_*PCs*_). The responses at time bin *m* were averaged across the remaining 3 trials for each exemplar, and assigned to a reference 2D matrix *R (28*_*exemplars*_*×12*_*PCs*_). Thus, every decoding iteration for training time point *m* vs. testing time point *n* began with a 2D test matrix *T*_*n*_ and reference, or training, 2D matrix *R*_*m*_: row *i* in matrix *T*_*n*_ is the vector of responses of all PCs to the randomly chosen single trial of the *i*^*th*^ exemplar at time *n*; row *i* in matrix *R*_*m*_ is the vector of averaged responses across the remaining 3 trials from all PCs, at time *m*, to the *i*^*th*^ exemplar. Next, we assigned 28 decoded labels based on the maximal correlation value between each row in the test and reference matrices: on every decoding step, the correlations between all rows of matrix *T*_*n*_ and all rows of matrix *R*_*m*_ were obtained, and the maximal value was detected. The label of the corresponding row in the reference matrix (e.g. “Face 1”) was then assigned to the row with the highest correlation to it from the test matrix. The assigned pair of test-reference rows were then excluded from subsequent steps in the current decoding iteration, such that every reference and test rows were assigned only once per iteration. Then, the next highest correlation value between the *T*_*n*_ and *R*_*m*_ rows was detected, and so on. After assigning all test-reference row pairs, the number of correctly assigned stimuli labels was recorded for that iteration. One thousand decoding iterations were performed for every training time bin *m* and testing time bin *n* combination, and the decoding accuracy for a specific *m* vs. *n* time bin combination was defined as the mean percentage of accurately decoded single exemplars across the 1,000 iterations. This resulted in a decoding accuracy matrix of size *28*_*training time bins*_ *× 28*_*testing time bins*_ in which each entry held the decoding accuracy for that specific training vs. testing time bins pair.

In order to obtain inter-temporal category decoding, we trained a k-nearest neighbors (k-NN) decoder (using k=1) on the PCA transformed data, iterating through all possible training vs. testing time bin combinations, similarly to the procedure in the single-exemplar decoding, described above. Classification accuracy was obtained via a leave-1-out validation method. Thus, for every training time bin *m* and testing time bin *n* (*m,n*= 1,2…28) combination, we trained the k-NN decoder on the category labels of 111 trials at time *m*, and tested its performance on the remaining trial at time *n* (28 different exemplars × 4 repetitions each = 112 trials in total). Category classification accuracy was defined as the percent of correctly decoded iterations, for every *m* vs. *n* time bins combination. This resulted in a *28*_*training time bins*_ *× 28*_*testing time bins*_ matrix, similarly to the exemplar decoding matrix, in which each entry detained the category decoding accuracy level for that specific training vs. testing time bins pair.

Statistical significance for the decoding analyses were assessed through random-shuffling of the item labels (single-exemplars/category labels, in accordance), across 1,000 permutations. In each shuffled permutation, the classifier accuracy scores we re-computed, as described above. P-values for each entry in the inter-temporal decoding accuracy matrices were defined as the proportion of shuffled accuracy scores larger than the actual classifier score. Correction for multiple comparisons was achieved via combined FDR correction (p=0.005) and cluster-size based correction (as detailed above in *Population activity patterns’ inter-temporal stability*).

