## Supplementary Data for "Perceptual stability reflected in neuronal pattern similarities in human visual cortex"

### Supplementary material

| Patient # | Sex | Age | Total electrode # | Visually responsive electrodes | Number of visually responsive electrodes in each ROI | | | |
|---|---|---|---|---|---|---|---|---|
| | | | | | Early visual (V1/V2) | Content-selective | Face-selective | Fronto-parietal |
| 1 | M | 28 | 146 | 30 | 4 | 14 | 9 | 2 |
| 2 | M | 50 | 106 | 36 | 7 | 12 | 0 | 1 |
| 3 | M | 38 | 102 | 24 | 0 | 16 | 1 | 2 |
| 4 | F | 44 | 160 | 26 | 4 | 10 | 4 | 2 |
| 5 | F | 29 | 128 | 43 | 9 | 12 | 4 | 0 |
| 6 | F | 22 | 241 | 12 | 0 | 0 | 0 | 11 |
| 7 | F | 34 | 146 | 28 | 4 | 6 | 4 | 2 |
| 8 | F | 30 | 188 | 21 | 0 | 9 | 4 | 1 |
| 9 | F | 29 | 287 | 53 | 0 | 10 | 2 | 25 |
| 10 | F | 32 | 219 | 9 | 0 | 0 | 0 | 4 |
| 11 | F | 55 | 270 | 18 | 1 | 3 | 2 | 1 |
| 12 | F | 27 | 385 | 42 | 3 | 17 | 10 | 3 |
| 13 | F | 33 | 193 | 35 | 0 | 5 | 3 | 12 |
| **Total** | | | **2571** | **377** | **32** | **114** | **43** | **66** |

**Table S1.** Details of patients and distribution of visually responsive electrodes across individuals.

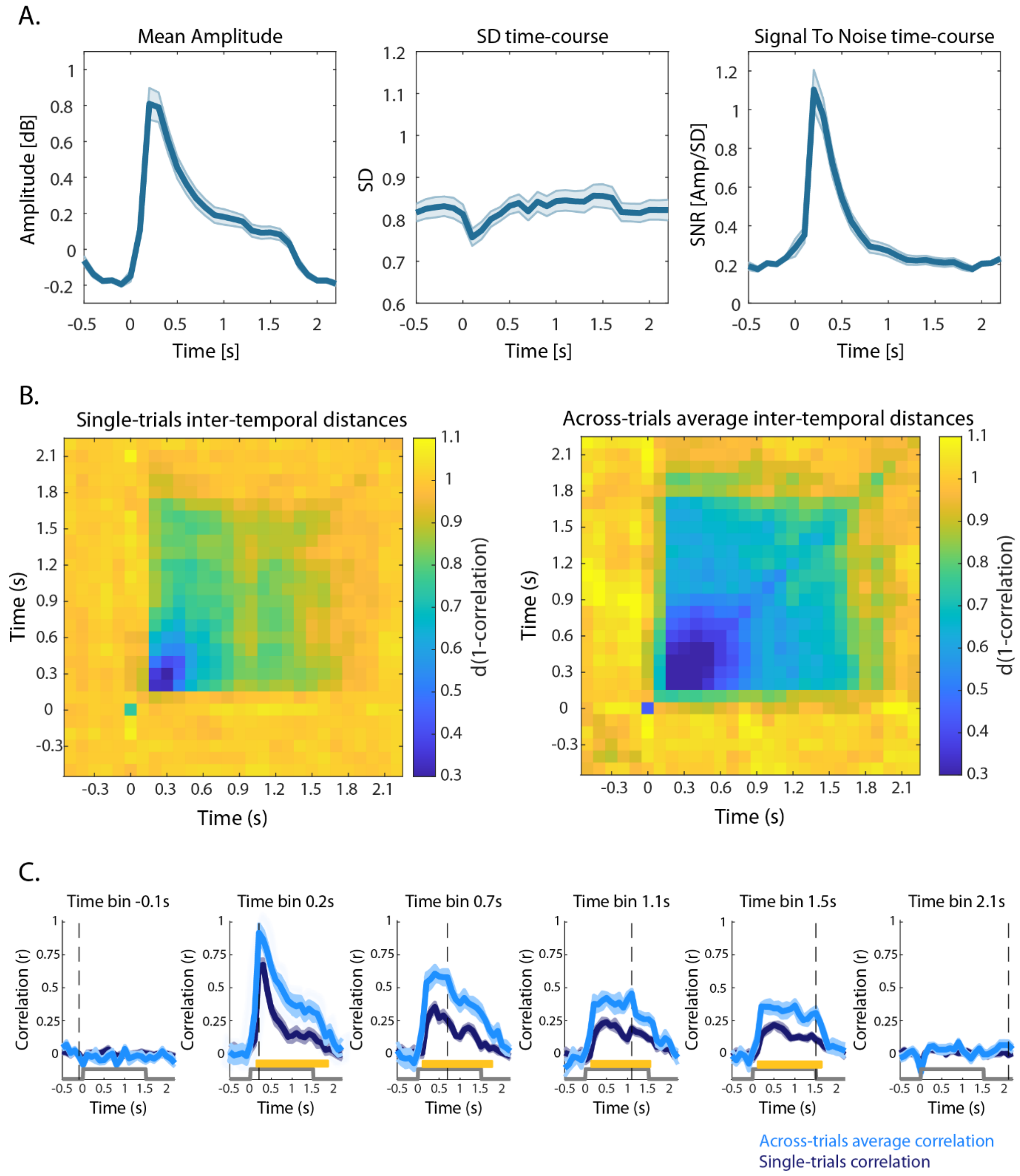


**Figure S1.** Signal-to-noise analysis and control in the content-selective electrodes (n=114). **A.** Mean amplitude, standard deviation and signal-to-noise ratio time-courses. **B.** Comparison of relational-coding inter-temporal distances matrices, derived from single-trial level correlations (left), vs. multi-trials average based-correlations (right), see *Methods* for further details. Notice how increasing the SNR through across- trial averaging resulted in higher, more sustained and persistent similarity levels (depicted as smaller distances, i.e. higher correlations, that were maintained across trial duration). **C.** Example relational coding inter-temporal correlation time-courses between 6 example time-bins, vs. all other time bins, for single-trial based correlations (dark blue), as compared to across-trials average-based correlations (light blue). Gray step-functions mark stimulus duration time. Yellow lines indicate significantly higher correlation levels for across-trials averaged data, as compared to correlations based on single-trials (paired t-test, $p<0.05$ fdr corrected).

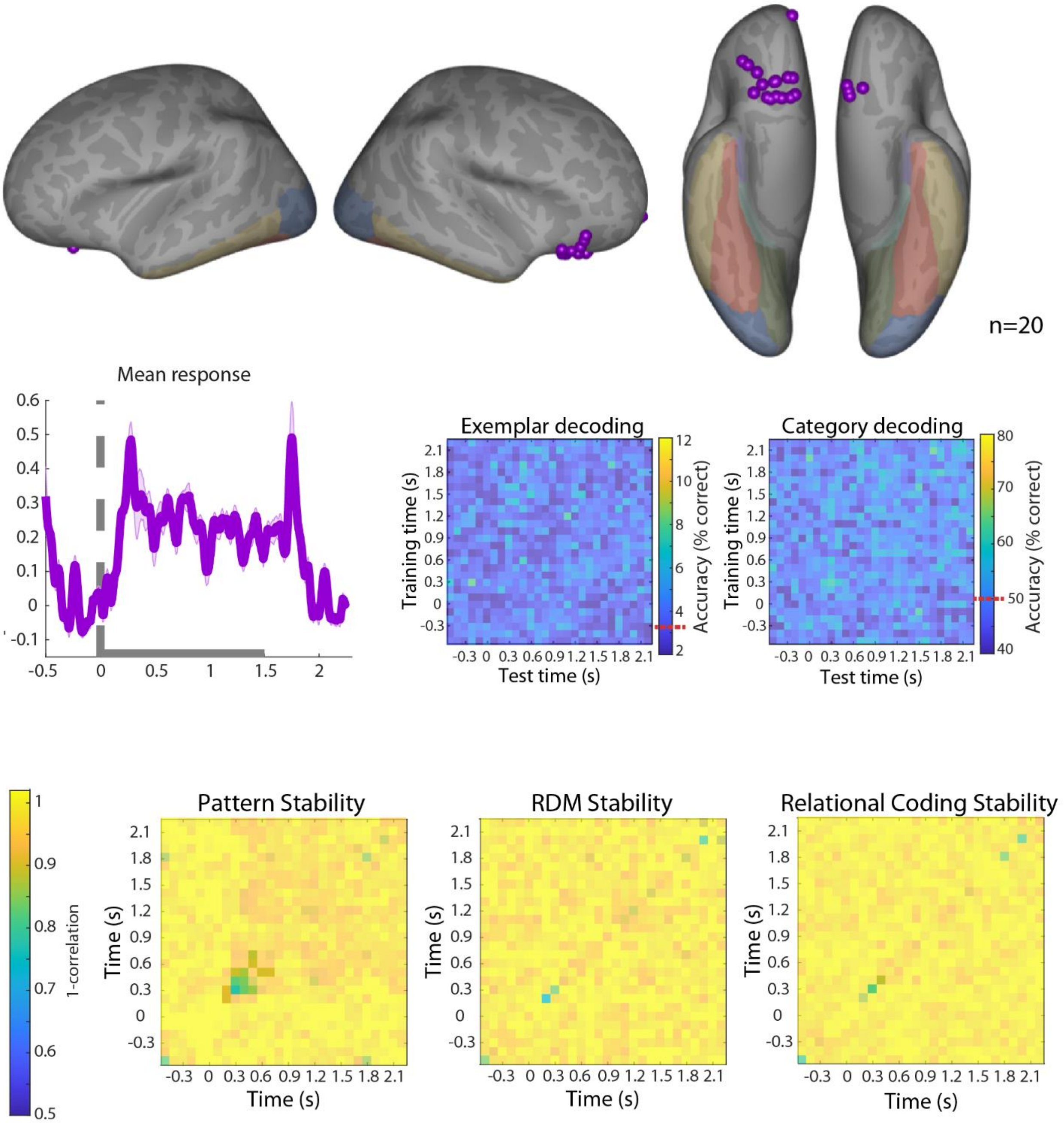


**Figure S2.** Results for visually-selective orbitofrontal cortex (OFC) electrodes (n=20): Top panels exhibit electrode locations on inflated cortical surfaces, from lateral (left) and ventral (right) views. Middle row panels include the mean response (similar to figure 2C), and exemplar and category inter-temporal decoding (similar to figure 7). Panels in bottom row display the inter-temporal distances matrices for pattern stability (similar to figure 4A), RDM stability (similar to figure 4C), and relational coding stability (similar to figure 5C). Half-transparent colored regions mark distances that are non-significant, as tested relative to distances emerged from random shuffling permutations, while distance values shown in opaque colors are statistically significant (1,000 random-shuffling permutations, p<0.01, fdr and cluster corrected).

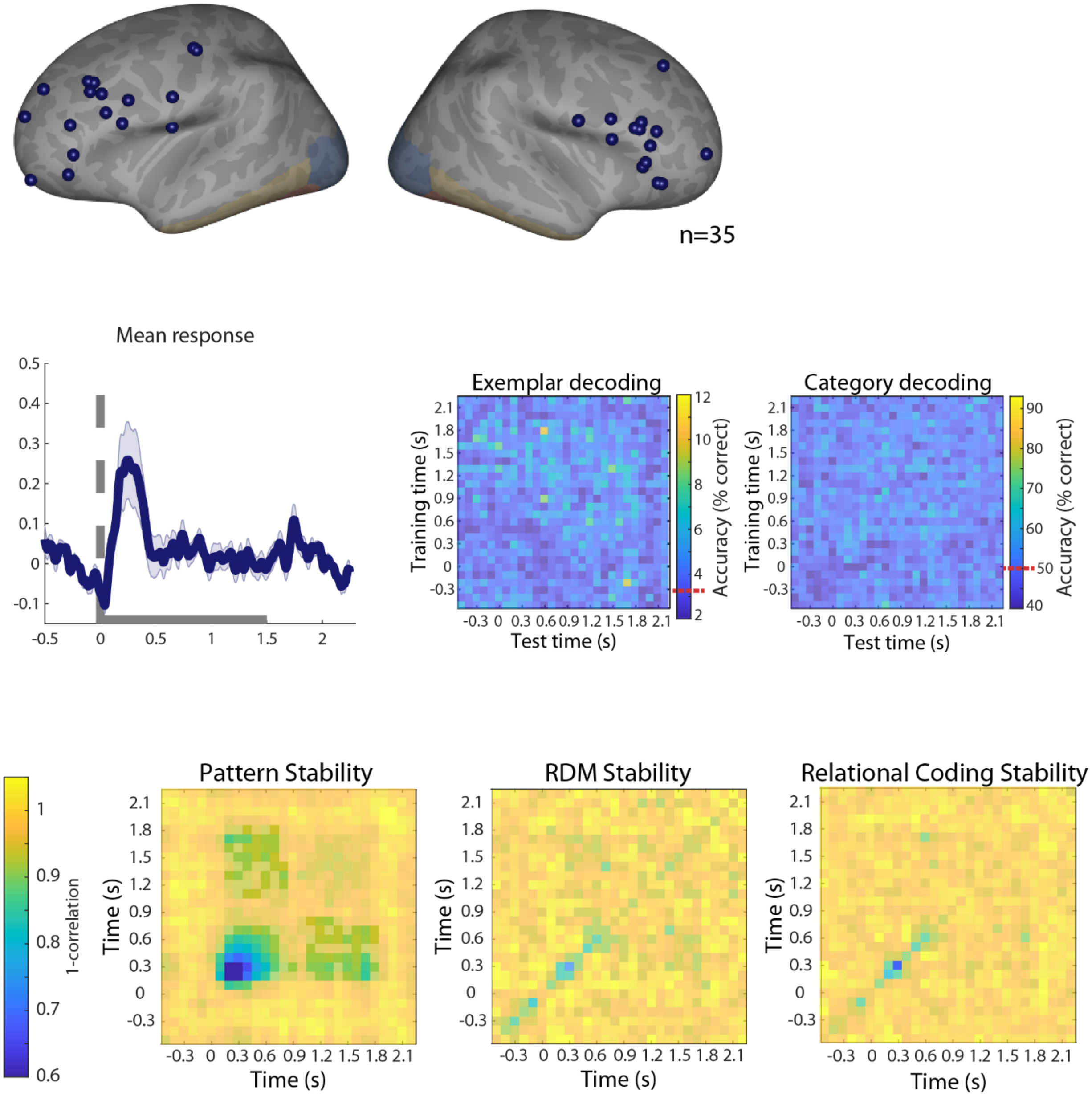


**Figure S3.** Results for visually-selective lateral-frontal electrodes (n=35), similar to as described in figure S2.

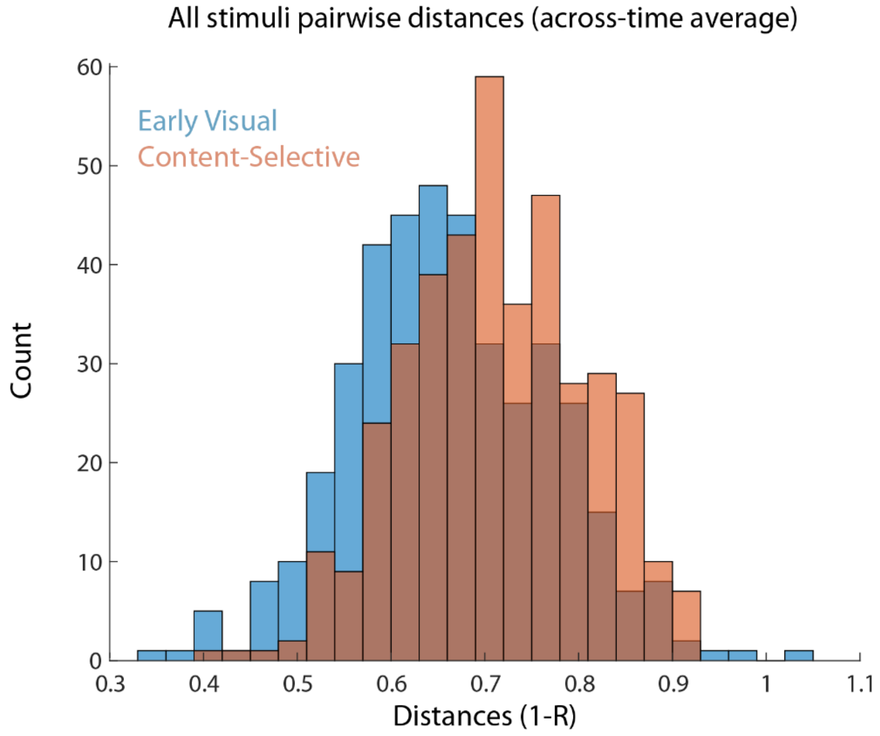


**Figure S4.** Histograms displaying all stimuli pair-wise distances (i.e. all the RDM values), defined as 1-R, calculated from the early visual cortex contact group (blue), and the high-order visual content-selective group (red). The RDM distances were calculated from the mean pattern activations at each 100ms time bin, and then averaged across time for the entire trial duration, and across all leave-1-out iterations of the same stimuli-pair (see *Methods*).

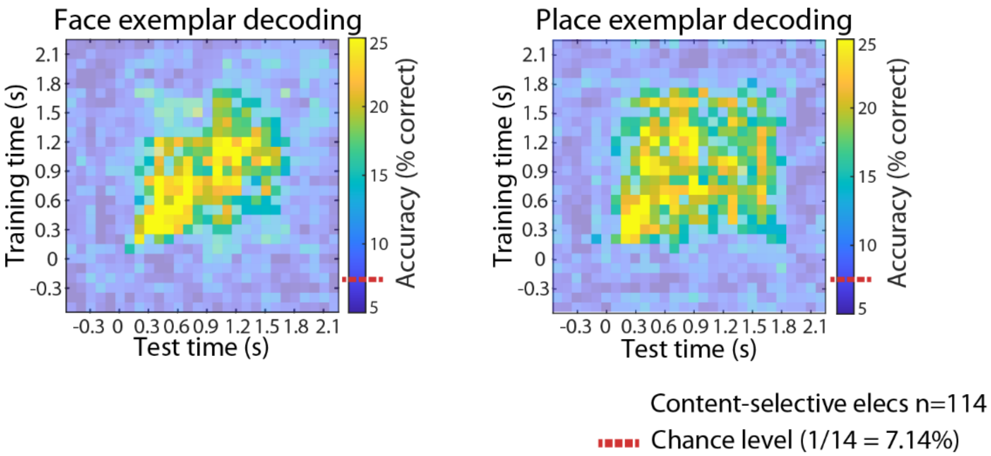


**Figure S5.** Inter-temporal exemplar decoding for each category separately, using content-selective contacts. An individual simple pattern-matching decoder was trained for each time bin, and then tested across all other time points. Dashed red lines mark chance level (1/14= 7.14%). Significant accuracy levels were calculated from a shuffling permutation test, comparing the real mean accuracy levels to the distribution of mean accuracy scores from 1,000 shuffled-labels permutations. Statistically significant decoding-accuracy levels are shown in opaque colors ($p<0.01$ fdr and cluster-based corrections), while insignificant values are shown as half-transparent. See *Methods* for further details.
